# AbrB antirepressor AbbA is a competitive inhibitor of AbrB-*phyC* interaction

**DOI:** 10.1101/2024.08.12.607668

**Authors:** Svetlana Neubauer, Manfred Roessle, Rainer Borriss, Oliwia Makarewicz

## Abstract

In *Bacillus* species, the interaction between the repressor AbrB and the antirepressor AbbA is vital for regulating gene expression. Our study reveals that AbrB binds DNA cooperatively the promoter of the *phyC* gene through multiple tetramers, forming a complex regulatory mechanism. AbbA disrupts AbrB’s DNA binding by competing for its DNA-binding sites, as shown by surface plasmon resonance (SPR) and gel shift assays. Circular dichroism (CD) confirmed that AbbA does not bind directly to the *phyC* promoter but mimics DNA to interfere with AbrB. Small-angle X-ray scattering (SAXS) data suggest that AbbA resembles a deformed DNA double helix. Our results indicate that AbbA binds to AbrB’s DNA-binding sites located with the N-terminal domain causing AbrB displacement. The interaction exhibits negative cooperativity, with both high- and low-affinity binding sites, as evidenced by Scatchard plots and kinetic studies. Our findings suggest that AbbA effectively mimics DNA to displace AbrB, activating transition-state genes. This research enhances our understanding of bacterial gene regulation and provides insights into the complex mechanisms controlling transcription in *Bacillus* species.

## Introduction

The transition from exponential bacterial growth to the stationary phase involves significant changes in gene expression to ensure survival under nutrient-limiting conditions. AbrB is a critical transition-state regulatory protein in *Bacillus species*, controlling the expression of over 100 genes that are upregulated post-exponentially and are involved in various cellular functions related to environmental changes. This tetrameric protein primarily acts as a repressor, with only a few genes known to be activated by AbrB [1–3].

AbrB-like proteins are found across almost all bacterial species and even in archaea, suggesting that this class of DNA-binding proteins is ubiquitous among prokaryotes. AbrB and related proteins, such as Abh and SpoVT, exhibit a ’swapped hairpin’ DNA-binding motif, related to double-ψ-β-barrels of MazE or VatN-N [4, 5] within their N-terminal regions. These proteins typically have a dimeric or tetrameric structure [6, 7].

The N-termini of AbrB/SpoVT-like proteins exhibit remarkably high sequence similarities (50-80%, see (the BLAST at ^1^), despite their interaction with different target genes and distinct cellular functions. In contrast, the C-terminal domain, which exhibits a domain-swapped fold, is thought to contribute less significantly to target gene recognition [8] and varies stronger between the related proteins.

An *in vivo* genome-wide analysis of AbrB and its paralogous protein Abh revealed that both proteins recognize hundreds of binding sites throughout the genome, with approximately 58% of these sites located within coding regions [9]. Among the 643 AbrB-binding sites identified, only 103 have been suggested to directly influence transcription. This has led to the controversial hypothesis that AbrB-like transcription factors may primarily function as nucleoid-associated proteins, similar to the histone-like proteins described in *E. coli* [10, 11].

At the transcriptional level, Spo0A, a key regulator of sporulation, is known to strongly repress *abrB* transcription [12]. Conversely, AbrB negatively auto-regulates its own promoter [13]. The highest levels of *abrB* transcripts are observed during the log phase and early exponential growth, with transcript levels nearly undetectable after the middle of exponential growth [14]. The protein levels of AbrB during growth align with these transcriptional profiles, with the effect primarily attributable to Spo0A [14].

In our previous studies, we demonstrated that AbrB represses phytase expression by binding to two distinct sites within the *phyC* region (ABS1 and ABS2) 15]. Surface plasmon resonance (SPR) experiments showed minimal dissociation of DNA from immobilized AbrB [15]. The equilibrium dissociation constants determined by SPR for binding to full-length AbrB were in the range of 10^-9^ M, indicating an unusually high binding affinity typically associated with antigen-antibody interactions [16–18]. Consequently, the low dissociation rate from the target gene would likely impede the bacterium’s ability to respond quickly to environmental stress.

Recently, a protein identified as an AbrB antirepressor, AbbA, was discovered [19]. Studies have shown that AbbA can replace or inhibit AbrB binding at the promoter regions of genes such as *sdp, skf, comK* and *lip*. Tucker A.T. and colleagues elucidated the high-resolution structure of AbbA using NMR spectroscopy [20, 21]. Each AbbA subunit is composed of three α-helices, with the second and third helices forming the intermolecular interface by interacting with their counterparts on the other subunit. Despite these insights, the precise mechanism by which AbbA binding inactivates the AbrB repressor remains unclear.

Here, we revisited the binding properties of AbrB and its antirepressor AbbA using surface plasmon resonance (SPR), experiments we initially conducted over a decade ago. Despite the significance of this research, the underlying mechanisms have not been published until now. Since then, the research group at HUB has disbanded, with the scientists moving on to new projects in different fields. Given the lack of progress in understanding the AbbA-AbrB interaction, we have decided to publish our research as a preprint on BioRxiv. We believe this data is valuable and may inspire further exploration within the scientific community.

## Material and Methods

### Bacterial strains, plasmids, and media

Bacterial strains and plasmids used in this study are listed in Table 1. Strains were grown in Luria Bertani (LB) medium with antibiotics added to the medium to final concentrations of 100 µg/ml ampicillin for AbrB and 50 µg/ml kanamycin sulfate for AbbA.

**Table 1.**
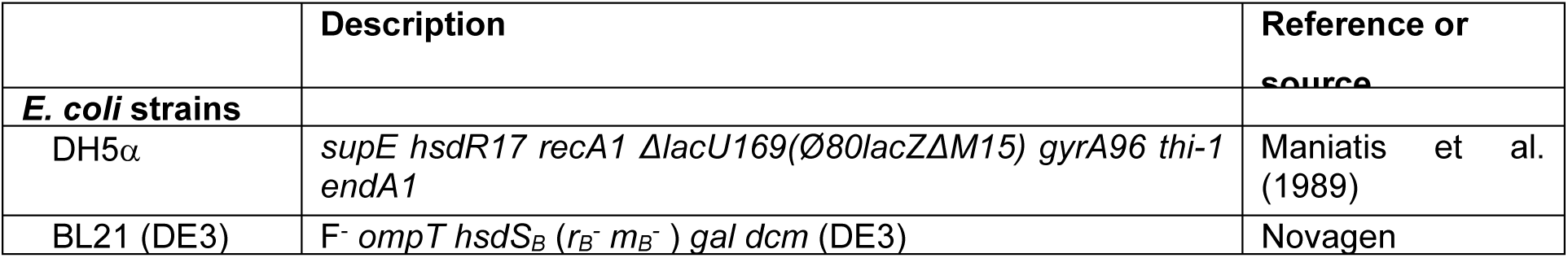
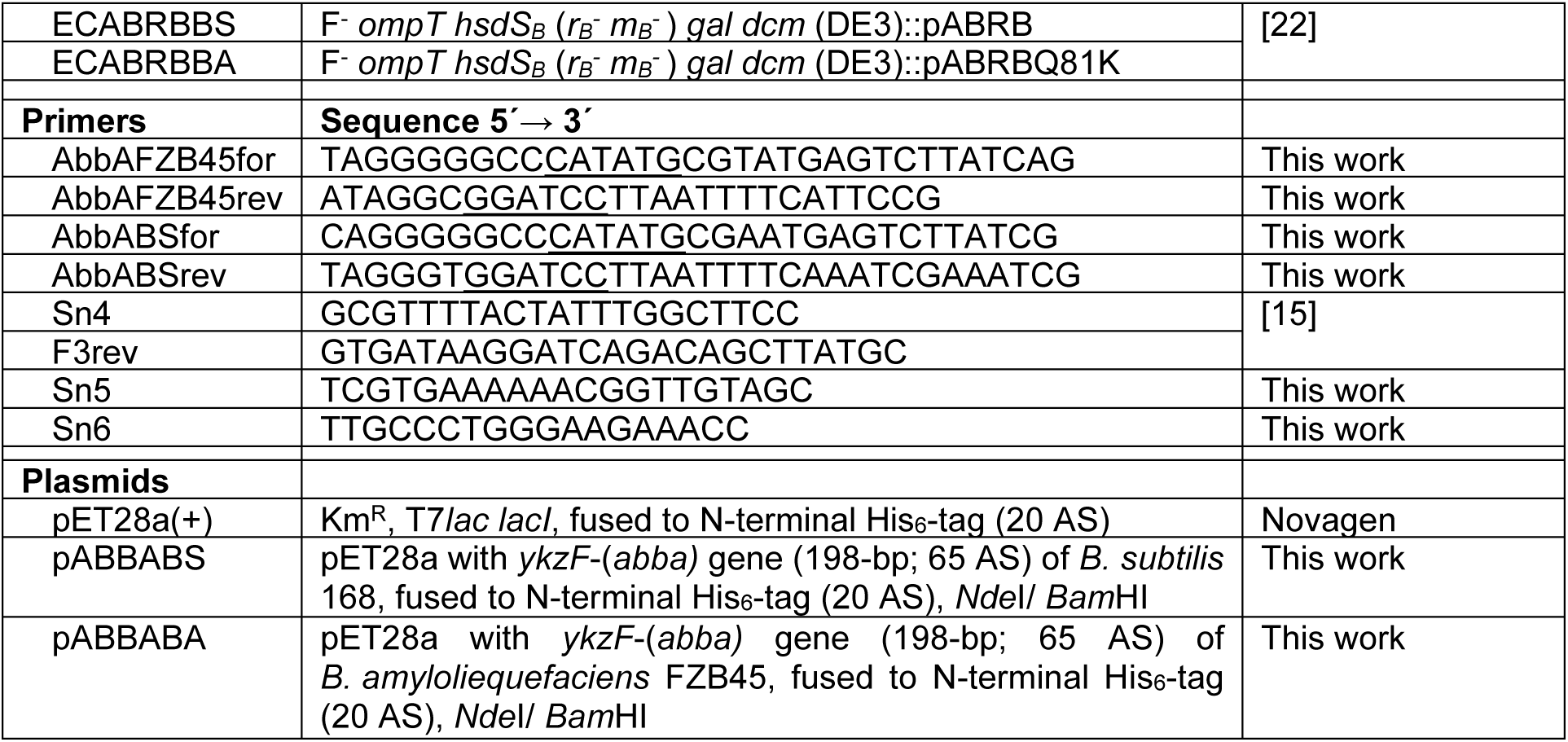
Strains, primers and plasmids used in this study.

### Construction of plasmids

The *abbA* genes were amplified from chromosomal DNA of *B. subtilis* 168 and *B. amyloliquefaciens* FZB45 using primers (Eurofins MWG Operon, Germany) listed in Table 1. PCR-products were purified using NucleoSpin®-Extract II kit (Machery-Nagel GmbH, Germany) and cloned into the *Nde*I and *Bam*HI cloning site of plasmid pET28a yielding plasmids pABBABS and pABBABA. *Nde*I and *Bam*HI digestions were done using enzymes purchased from Fermentas GmbH (Germany) according to manufacture’s instructions. The plasmids were transformed into Ca^2+^ competent DH5α cells and transformants were selected on LB-agar plates containing 50 µg/ml kanamycin. Plasmids were isolated using NucleoSpin®-Plasmid kit (Machery-Nagel GmbH, Germany), conformed by sequencing (SMB, Berlin, Germany) and transformed into Ca^2+^ competent *E. coli* BL21 cells.

### Overexpression and purification of the proteins

*E. coli* BL21 cells bearing the expression plasmids were grown in LB containing 1 % lactose by shaking at 200 rpm and at 37 °C overnight. The cells were harvested in a Sorvall RC5B centrifuge (Thermo Scientific) at 5000 rpm (in a GS3 rotor), washed with 100 ml P-buffer (50 mM Tris HCl pH 7.4, 300 mM NaCl, 10 % glycerol, 1 mM DTT for AbbA and 2 mM β-mercaptoethanol for AbrB). The cell pellet was resuspended in 10 ml P-buffer and incubated with 1 mM PMSF (Serva) and 25 U benzonase (Novagen) on ice for 20 min. Lysis of cells was carried out by sonification. The proteins were bound on Ni^2+^-NTA agarose (Qiagen) and incubated at 4°C for 2 h under agitaion. After extensive washing with P-buffer (40 ml), and P-buffer containing 30 mM, 50 mM and 100 mM imidazol the His-tagged AbrB proteins were eluted with 350 mM imidazol and His-tagged AbbA proteins with 450 mM imidazol in P-buffer in 500 µL fractions. Eluted fractions were analysed in 14% SDS-PAGE and pure fractions were pooled. N-terminal His-tags of both AbbA proteins were removed using Thrombin CleanCleave^TM^ Kit (Sigma-Aldrich, Germany) according to the manufacture protocol. After thrombin cleavage, the resulting protein contained 3 additional (GSH) N-terminal residues. Proteins were dialysed stepwise against buffers containing 25 % glycerol and 50 % glycerol. Dialysis buffer conditions differ in dependence of the experiments and proteins: For gel shift assays 50 mM Tris HCl (pH 7.4), 300 mM NaCl, 2 mM β-mercaptoethanol (AbrB) or 1 mM DTT (AbbA) and 50 % glycerol were used. For measurement of circular dichroism (CD) 10 mM Tris HCl (pH 7.4), 300 mM NaF, 10 mM DTT (AbrB) or 1 mM DTT (AbbA) and 10 % glycerol were used. For surface plasmon resonance measurements 10 mM HEPES (pH 7.4), 300 mM NaCl, 10 mM DTT (AbrB) or 1 mM DTT (AbbA) and 10 % glycerol were used. For small-angle X-ray scattering, the proteins were dialysed in 50 mM Tris HCl (pH 7.4), 150 mM NaCl, 2 mM β-mercaptoethanol (AbrB) or 1 mM DTT (AbbA). Protein concentrations were determined at 280 nm using NanoDrop ND2000 (Peqlab Biotechnologie).

### FPLC

For CD spectroscopy and SAXS measurement (see below) proteins were additionally purified using gel filtration chromatography (ÄKTA purifier, GE Healthcare) on a semi-preparative Superdex™ 200 column (GE Healthcare) using buffer conditions mentioned above. Protein samples (300 µl – 500 µl) were applied on the column and run with the flow rate of 0.2 ml / min at room temperature. Protein content of the flow through was measured at 280 nm. Fractions of 200 µl were collected and analysed by 14% SDS-PAGE. The column calibration was performed by running a mixture of standard proteins with known molecular masses: vitamin B12 (1.35 kDa), myoglobin (17 kDa), ovalbumin (44 kDa) und alpha-globulin (158 kDa).

### Native agarose electrophoresis

Complex formation between AbrB and AbbA variants was analysed under native conditions using 2 % agarose gels prepared and run in 1xTG buffer (25 mM Tris HCl pH 8.8, 192 mM glycin). These assays were carried out using purified His_6_-AbbA and purified His_6_-AbrB. The His_6_-Tag of the both proteins has no effect on *in vitro* binding assays as described previously [19]. The isoelectric points (IEP) of the proteins have been calculated with IEP 6.24 for AbbA_BS_ and IEP 6.15 for AbbA_BA_. Since AbrB and AbrBN exhibit high isoelectric points (IEP of 8.3 for AbrB_BS_, IEP of 8.9 for AbrB_BA_ and IEP of 9.3 for AbrBN) free proteins were seen to stick in the slots or run to cathode in previous experiments using native gels. Therefore, the slots were prepared in the middle of the gels and 10 µl agarose was applied to close the slots after loading of the samples. Gels were run at room temperature for 90 min at 70 V, then gels were rinsed in fixing-solution (20 % ethanol and 10 % acidic acid) for 10 min and washed in deionised water 4 times for 30 min. Gels were placed on glass plates, wrapped in wet thin filter papers and dried at 37°C overnight. Proteins were dyed using PageBlue™ Protein Staining Solution (Fermentas) and digitalized using the Molecular Imager FX-Pro Plus (BioRad).

### Gel shift assay

A 440 bp DNA fragment corresponding to the *phyC* region (−333 to +107) was amplified using primers Sn5 and Sn6 listed in table 1 and purified using NucleoSpin®-Extract II kit (Machery Nagel). Gel shift assays (EMSA) were carried out using purified His_6_-AbbA and purified His_6_-AbrB. For gel shift assays the fragment was 5′-^32^P labelled by [γ-^32^P]-ATP (Hartmann analytic) using T4 polynucleotide kinase (PNK) (Fermentas) and purified using QIAquick PCR Purification kit (Qiagen) according to manufactures’ protocols, respectively. Gel shift assay experiments were performed as described previously [22]. The binding reaction was carried out with 10000 cpm of DNA (corresponding to 1.14 nM). A labelled 523 bp DNA fragment (−392 to +131, Sn4/ F3rev-PCR product, 10000 cpm corresponding to 0.37 nM) was used for control assay with various AbbA concentrations in absence of AbrB.

### Surface plasmon resonance measurement

Immobilization of the AbrB proteins (ligand) on CM5 chips (GE Healthcare) and interaction analyses were performed as described previously [15]. AbrB proteins to be covalently bound to the matrix were diluted in 50 mM HEPES (pH 7.4), 300 mM NaCl, 10 mM DTT and 10 % glycerol to a final concentration of 100 nM and 10 µl were applied on the activated chip at a flow rate of 5 µl/min. In total 470 response units (RU) of full-length AbrB_BA_ (9.29 fmol/mm^2^ corresponding to a tetramer) were coupled at flow cell 4 (Fc4), 484 RU of full-length AbrB_BS_ (9.57 fmol/mm^2^ corresponding to a tetramer) at Fc3 and 143.6 RU of the truncated AbrBN at Fc2 (9.49 fmol/mm^2^ corresponding to a dimer).

The AbbA proteins (analyte) were diluted in 50 mM HEPES (pH 7.4), 300 mM NaCl, 1 mM DTT and 10 % glycerol and sequentially injected over the immobilized proteins at different concentrations (0,05 – 5 µM) at a constant flow of 60 µl/min, with binding being monitored for 135 seconds, after the end of injection followed by a 115-seconds dissociation period. The uncoupled Fc1 was used as reference and was subtracted from all raw data.

The SPR sensorgrams were evaluated by BIAevaluation 3.1 software package using numerical integration algorithms representing different binding models. These include simple bimolecular (Langmuir) binding (A+B ↔ AB), conformational change (A+B ↔ [AB] ↔ A×B), bivalent analyte (A+B ↔ AB+B ↔ AB2), and additional a binding model of heterogeneous analyte (competing reactions: A1+B ↔ A1B; A2+B ↔ A2B) and heterogeneous ligand (parallel reactions: A+B1 ↔ AB1; A+B2 ↔AB2). The best fitting was determined by analysing the χ^2^ (χ^2^<1% of RU_max_) at the steady-state.

Since no satisfactory results for global fitting of the association and dissociation constants could be obtained only the thermodynamical parameters of the steady-state plots were analyzed as described previously [15]. Each SPR-curve could be successful fitted separately (local fit) using definite binding models to obtain the kinetic parameters. Hill-coefficients were derived from Hill-plots at steady as described previously [15].

### Circular dichroism measurement

Circular dichroism measurements were performed on a Jasco J-720 spectropolarimeter in 0.1 mm (Starna) or 1 mm (Hellma) quartz cuvettes. CD-spectra were recorded between 185 nm and 320 nm. Five scans for DNA-protein or protein-protein interaction and twenty scans for analysis of secondary structure were accumulated and averaged. We used a the D4-ds-oligonucleotide as DNA target derived from AbrB-binding site 2 of *phyC* promoter region that was shown to exhibite high binding affinity to AbrB [23]. The buffer reference spectra were used as baselines and were subtracted from the CD-spectra of the samples. To avoid any effects of the slight variance in test conditions or instrumentation, every day new buffer and protein references were run. The binding studies were performed at concentrations of 20 µM AbrB, 40 µM AbbA and 2.5 µM oligonucleotide, respectively. Since AbrB is a homotetramer in solution [24] the used concentration corresponded to 5 µM AbrB_4_. The spectra of oligonucleotides were subtracted from the spectra of proteins to visualize the changes of the proteins. Signal intensities were expressed in mean molar ellipticity per residue [Θ]_MRE_ (deg cm^2^ dmol ^-1^ residue^−1^) according to the equation (1)

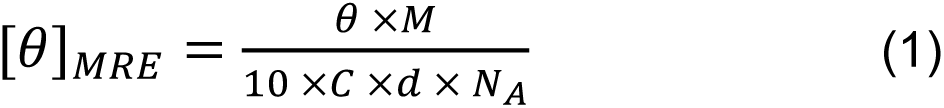

where ‘NA’ represents the number of amino acid residues, ‘d’ is the path length of the cell [cm], ‘c’ is the concentration [g/L] and M is the molecular mass [Da].

The analysis of protein secondary structures was performed on DichroWeb server [25] using SELCON3 [26–28], CONTIN [29, 30], CDSSTR [28, 31] algorithms and different reference datasets. As CD data were only reliable above 185 nm, datasets 3, 4, 6, 7 and SP175 were used. The normalized root mean square deviation (NRMSD) was used as a goodness-of-fit parameter of each deconvolution output [25].

### Small angle X-ray scattering

After Ni^2+^-NTA agarose purification (see above) the proteins were additionally purified by Protino Ni-TED 1000 (Macherey-Nagel) followed by FPLC gel filtration (see above). After purification and dialysis, the protein were concentrated by using Vivaspin® centrifugal filters (MMCO 15 kDa and 30 kDa).

SAXS data were collected for 2.5 mg/ml and 5 mg/ml of each protein in dialysis buffer following standard procedures at beamline X33 at the European Molecular Biology Laboratory (EMBL) on the DORIS III storage ring (DESY, Hamburg, Germany). The momentum transfer range of s = 0.08-5.0 nm^-1^ (s = 4π sin(θ)/λ, 2θ representing the scattering angle of 0.1 to 10°, λ = 1.5 Å) was covered at a sample-detector distance of 2.7 m using exposure time of 2 min at 8°C [32, 33]. The sample volumes were adjusted at 60 µl.

The scattering patterns were visualized as radially averaged one-dimensional curves. From these curves several important overall parameters can be directly obtained providing information about the size, oligomeric state and overall shape of the molecule [34]. SAXS data analysis was performed using the software suite ATSAS [35, 36] including algorithms PRIMUS, GNOM, GASBOR, DAMAVER. PRIMUS was applied to manipulate the experimental data files like averaging of scattering of signals over all orientations due to isotropic intensity, subtraction of solvent scattering pattern, and merging of data sets obtained for both protein concentrations. The final curve (SM6) was fitted and evaluated providing the integral parameters from Guinier plots [37] such as radius of gyration R_g_ (for globular, flat and rod-type particles). The pair distance distribution functions (P(r)) were evaluated using the indirect transformation algorithm GNOM [38, 39], yielding also the maximum particle diameter (D_max_).

*Ab inition* modeling was performed using the algorithm GASBOR that reconstructs protein structure by a chain-like ensemble of dummy residues [33, 40]. Proteins typically consist of folded polypeptide chains composed of amino acid residues separated by 0.38 nm between adjacent C_α_ atoms in the primary sequence. At a resolution of 0.5 nm, a protein structure can be considered as an assembly of dummy residues (DR) centred at the C_α_ positions. The number of DRs is equal to the number of residues in the protein sequence. A dummy solvent atom was placed 0.5 nm outside the protein. GASBOR processes the output files of GNOM including extracted parameters (such as R_g_, D_max_), particle symmetry (P22 for AbrB, P2 for AbbA) and the number of residues within a single asymmetric unit. DAMAVER superposed the fifteen (for AbbA) or twenty (for AbrB) individual evaluated models and subsequently averaged these models which were aligned to yield the best overlap. DAMAVER generated a list of models as PDB files which were visualized by PyMol and RasMol software.

## Results and Discussion

### Analysis of the polymerization state of the proteins

The quality and polymerization states of all proteins were assessed using analytical gel-filtration chromatography (FPLC). It has been previously reported that active AbrB from *B. subtilis* (AbrB_BS_) forms tetramers in solution [24]. AbrB from *B. amyloliquefaciens* FZB45 (AbrB_BA_) differs from AbrBBS by only a single amino acid residue (Q81K), resulting in 98.9% identical amino acid residues. Both proteins have shown similar binding properties to the phyC gene region [22]. Therefore, similar polymerization states were expected and were confirmed experimentally. Both AbrB variants eluted as single peaks at volumes corresponding to their tetrameric states (Figure 1A).

**Figure 1.**
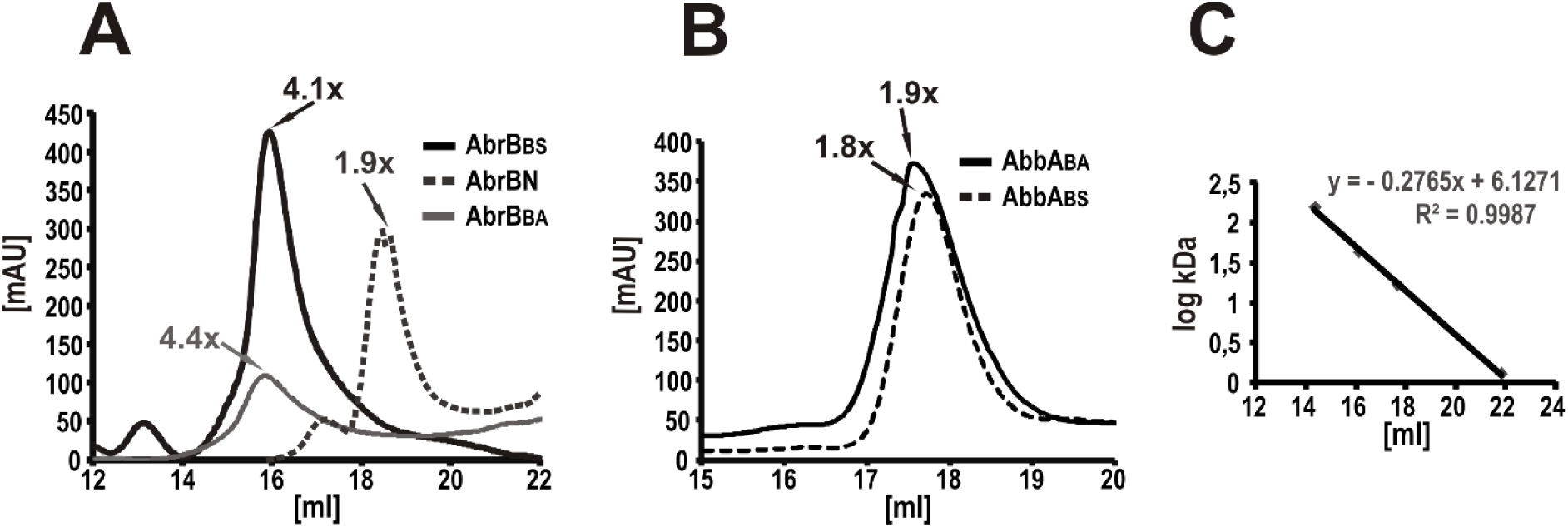
FPLC of the AbrB (A) and AbbA (B) variants of *B. subtilis* 168 and *B. amyloliquefaciens* FZB45. Preparativ filtration was performed on a Superdex™ 200 column (GE Healthcare). Absorbance [mAU] at 280 nm was plotted versus the elution volume [ml]. (C) The column calibration was performed by running a mixture of standard proteins (plot of logarithm of the known molecular masses versus the elution volume).

The AbrB antirepressor, AbbA, was first described in 2008 [19]. Recent studies using size-exclusion liquid chromatography (SELC) have identified AbbA from *B. subtilis* as a dimer [21]. In our experiments, we analyzed both AbbA variants, AbbA_BS_ and AbbA_BA_, using analytical FPLC. Both proteins were eluted as single peaks corresponding to their dimeric states (Figure 1B).

We performed an alignment (using ClustalW) of the amino acid sequences of AbbA from various *Bacillus* species (Figure 2). The alignment revealed 22 identical amino acids, including negatively charged glutamate residues (E12-14, E30, E35, E40) and aspartate residues (D38, D47), with fewer positively charged residues (K44, K52, R60). Additionally, highly conserved residues include hydrophobic residues with branched chains (L17-19, L22, L32, L57, and I39), small residues (A28, G42), and aromatic residues (Y27, Y51).

**Figure 2.**
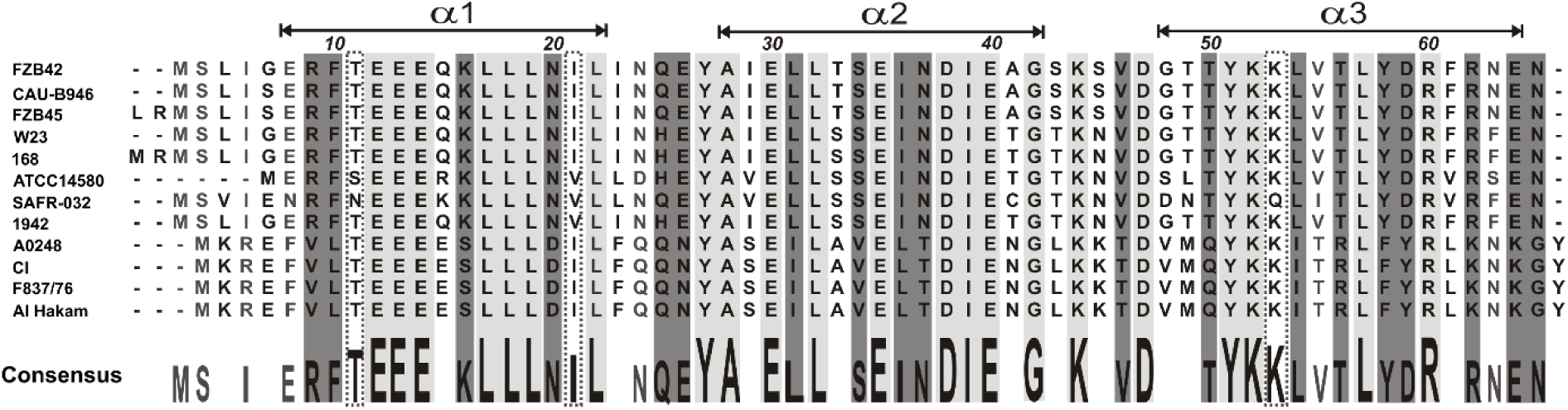
Amino acid sequence alignment of AbbA-like proteins of *Bacillus* related species. FZB42 = *Bacillus amyloliquefaciens* FZB42; CAU-B946 = *Bacillus amyloliquefaciens* subsp. *plantarum* CAU-B946; W23 = *Bacillus subtilis* subsp. *spizizenii* W23; 168 = *Bacillus subtilis* subsp. *subtilis* 168; ATCC14580 = *Bacillus licheniformis* ATCC14580; SAFR-032 = *Bacillus pumilus* SAFR-032; 1942 = *Bacillus atrophaeus* 1942. The sequence of AbbA from *Bacillus amyloliquefaciens* FZB45 was sequenced within this work (SMB, Berlin). The pathogenic Bacillus species: A0248 = *Bacillus anthracis* A0248; Cl = *Bacillus cereus* biovar. *anthracis* Cl; F837/76 = *Bacillus cereus* F837/76; Al Hakam = *Bacillus thuringiensis* Al Hakam. Structural regions (α1-3) are indicated on the top according to Tucker 2013 [21]. The consensus sequence obtained from this alignment is given on the bottom. Identical residues are represented as a upper case on a light grey background. Corresponding residues with 100 % < AS > 75 % (Lys53, Thr11, Ile21) are shown in dotted boxes. The moderate conserved residues of 66 % similarity are represented on a dark grey background; the residues with 58 % similarity are highlighted in grey. The sequence alignment was performed with ClustalW (DS Gene 1.5).

The AbbA proteins from *B. subtilis* and *B. amyloliquefaciens* FZB45 are highly conserved, sharing approximately 89% of identical amino acid residues. This conservation is notable but reveals somewhat greater variation compared to the AbrB variants.

### Analysis of the formation of protein complexes

We monitored the complex formation between AbrB and AbbA using native agarose gel electrophoresis. The His6-tagged AbrB proteins exhibit high isoelectric points (IEPs): AbrB_BS_ showed an IEP of 8.3, AbrBBA showed an IEP of 8.9, and AbrBN showed an IEP of 9.3, as theoretically calculated by DSGene 1.5 software. In contrast, the AbbA proteins had lower IEPs, with AbbA_BS_ at 5.04 and AbbA_BA_ at 4.88. Notably, His6-tagged AbbA proteins, which were used for binding studies, exhibited slightly higher IEPs (details in Materials and Methods).

Due to these differences in IEPs, AbrB and AbbA migrate differently under electrophoretic conditions and a common native PAGE was not suitable for this analysis. Instead, we used horizontal agarose gels at pH 8.8, where samples were applied in the center of the gel. This setup allowed free proteins to migrate according to their IEPs, while the formed complexes migrated in the direction corresponding to their combined IEP.

As anticipated, the migration behaviors of the free AbrB proteins varied: AbrB_BS_ migrated slowly toward the anode (+), AbrB_BA_ remained stacked in the slots at pH 8.8, and the truncated AbrBN migrated toward the cathode (−). Both AbbA proteins migrated toward the anode.

Binding assays were conducted using constant concentrations of AbrB (10 µM for AbrB and 15 µM for AbrBN) and varying concentrations of AbbA (5 µM, 10 µM, and 20 µM for AbrB; 7.5 µM, 15 µM, and 30 µM for AbrBN) and vice versa. The concentration-dependent complex formation was assessed for the corresponding AbrB and AbbA proteins. The electrophoretic mobility of full-length AbrB changed in the presence of AbbA, and vice versa. Stable AbrB-AbbA complexes from both *Bacillus* species were observed (Figure 3A and B) without smearing bands, indicating well-defined interactions.

**Figure 3.**
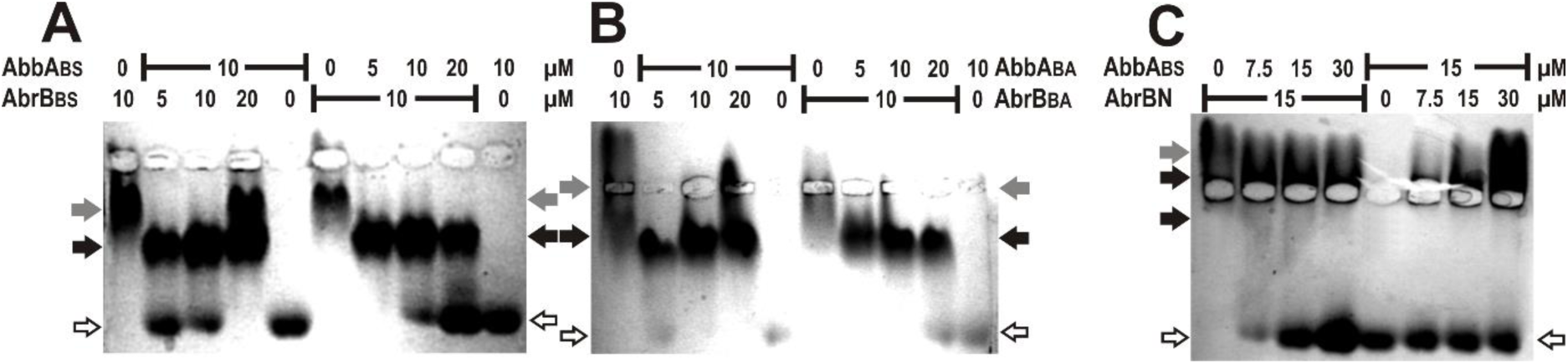
Native agarose electrophoresis for analysis of the complex formation of AbrB and AbbA variants. The concentration of AbrB (10 µM) or AbrBN (15 µM) was kept constant and the AbbA proteins from *B. subtilis* 168 (BS) and *B. amyloliquefaciens* FZB45 varied and conversely. White arrowheads, unbound AbbA; grey arrowheads, unbound AbrB; black arrowheads, AbrB-AbbA-complex.

At increasing AbrB concentrations and constant AbbA concentration, an AbrB band appeared suggesting saturation of AbbA with AbrB. A similar effect was noted at higher AbbA concentrations with a constant AbrB concentration, where a free AbbA band was visible. The saturation was achieved at a stoichiometry of approximately 1 AbrB-tetramer : 2 AbbA-dimers. In contrast, complex formation between the truncated AbrBN and AbbA_BS_ was relatively weak. At higher AbbA concentrations, AbrBN exhibited a slight change in mobility, with the AbrBN band shifting closer to the slots, suggesting a slower migration toward the cathode.

We also introduced an alanine substitution at the highly conserved N-terminal arginine residue at position 15 (R15A) of AbrB_BS_, which is thought to play a crucial role in DNA interactions [41]. In previous studies, we showed that substitution of arginine R15 to alanine results in an increase of octameric forms of AbrB but contrary to expectations, this mutant did not completely lose its ability to bind the 440 bp *phyC*-DNA fragment [42]. However, the R15A mutant notably lost its capacity to form a complex with the antirepressor AbbA (Figure 4A).

**Figure 4.**
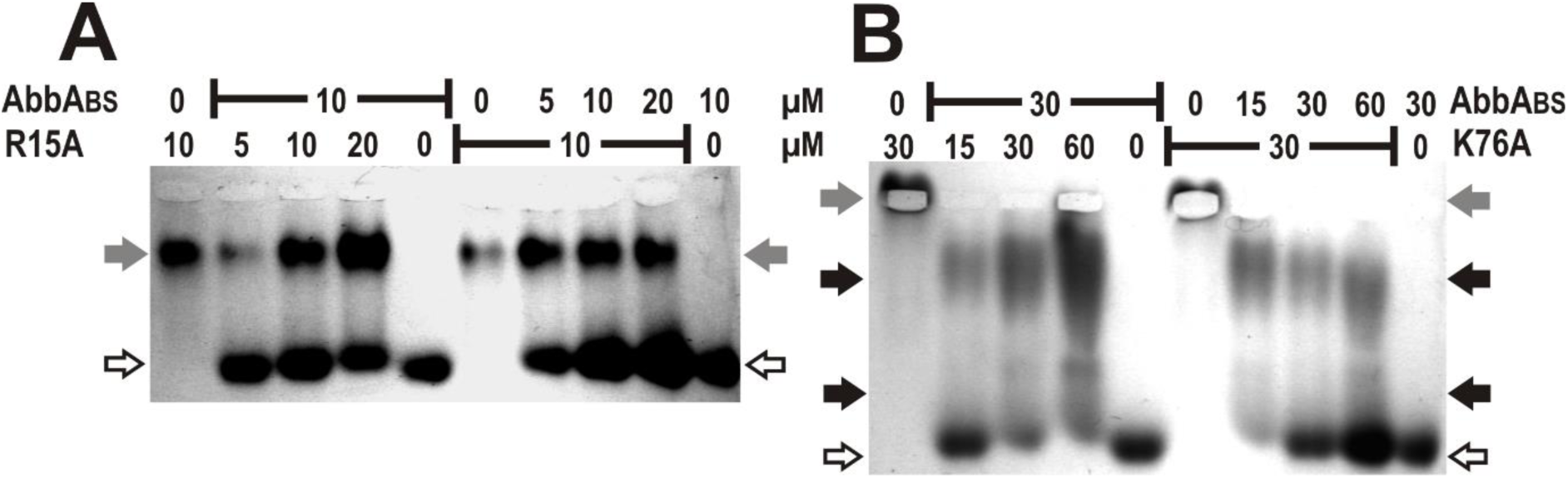
Native agarose electrophoresis for analysis of the complex formation of AbrB and AbbA variants. The alanine substitution of the arginine residue at position 15 (R15A) was performed within the N-terminal DNA-binding domain and lysine exchange (K76A) within the C-terminal domain. White arrowheads, unbound AbbA; grey arrowheads, unbound AbrB; black arrowheads, AbrB-AbbA-complex.

Additionally, we constructed a mutant of the C-terminal domain of AbrB_BS_ by substituting lysine 76 with alanine (K76A), which resulted in reduced DNA-binding properties [42]. When tested in the presence of AbbA, the mobility of the AbrBK76A mutant changed (Figure 4B). The interaction between AbrBK76A and AbbA formed an unstable complex, as evidenced by the smearing bands observed in the gel.

### Effect of AbbA on phyC repression by AbrB

It was previously reported that AbbA effectively blocks AbrB binding to the promoters of various genes, including promoters of *sdp*, *skf*, *lip* and *comK* in gel retardation experiments [19]. In our earlier studies, we demonstrated that both AbrB proteins AbrB_BS_ and AbrB_BA_ can shift the promoter region of the *phyC* gene in a concentration-dependent manner [22].

Here, we analyzed the ability of AbbA_BS_ and AbbA_BA_ to block interaction of AbrB_BS_ and AbrB_BA_ to a 440 bp *phyC*-fragment (−333 to +107) using gel retardation assays (Figure 5). In these experiments, we maintained constant concentrations of 1 µM AbrB_BS_ and 2 µM AbrB_BA_, while varying the concentrations of AbbA (ranging from 0.5 µM to 6 µM). Notably, AbbA proteins alone were unable to shift the *phyC*-DNA (Supplementary Material (SM) 1).

**Figure 5.**
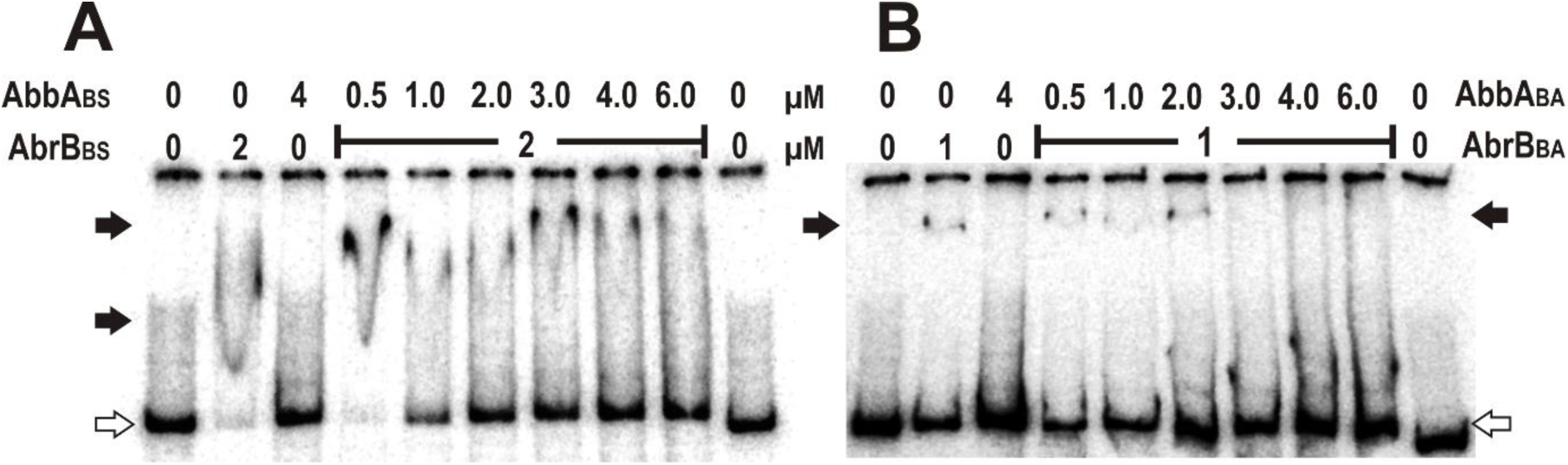
Gel shift assays with AbrB and AbbA proteins in presence of the *phyC* region and increasing concentrations of AbbA. The control lanes (0 µM) are without AbrB or AbbA protein as well as without both proteins. The 440 bp DNA fragment, harboring the entire *phyC* promoter region with AbrB binding sites was added at constant 1.14 nM concentration (10000 cpm/µl). White arrowheads, free DNA; black arrowheads, AbrB-DNA complex.

When 2 µM AbrB_BS_ was combined with 0.5 µM AbbA_BS_, no free DNA was observed. However, at higher AbbA_BS_ concentrations (from 1 µM to 6 µM), the AbrB_BS_-DNA complex began to dissociate in a stepwise manner, leading to the release of the DNA. This stepwise dissociation was consistent with previous observations made during gel shift experiments with the full-length *phyC* fragment and increasing AbrB concentrations in the absence of AbbA, which also resulted in smearing bands at concentrations up to 1 µM AbbA_BS_.

Interestingly, the two AbrB proteins exhibited differences in how they dissociated from the *phyC* gene region when exposed to their respective AbbA proteins. At 3 µM AbbA_BA_, the AbrB-DNA complex was no longer observed, and the dissociation of the DNA-AbrB_BA_ complex occurred rapidly and without intermediate forms. In both cases, at least a fourfold excess of AbbA dimers was required to fully dissociate the AbrB-DNA complex.

These results clearly demonstrate that AbbA from both *Bacillus* species inhibits the ability of AbrB to bind to *phyC*-DNA, thereby acting as an effective antirepressor.

### Evaluation of binding kinetics from the SPR data

To investigate the binding kinetics and thermodynamic properties of soluble AbbA with immobilized AbrB, we utilized surface plasmon resonance (SPR), a sensitive label-free optical biosensor technique. Representative sensorgrams illustrating the interactions between AbbA and AbrB from *B. subtilis* and *B. amyloliquefaciens* FZB45, as well as the truncated AbrB protein (AbrBN), are presented in SM 2.

Our binding studies revealed that the interaction between soluble AbbA and immobilized AbrB is concentration-dependent across a range of 0.05 µM to 4 µM. During the association phase, the instrument response increased in direct proportion to AbbA concentration, clearly demonstrating a concentration-dependent binding relationship.

Analysis of the binding curves for the interaction between soluble AbbA and immobilized full-length AbrB_BS_ and AbrB_BA_ proteins, performed using BIAevaluation 3.1 software, revealed that the data could be described by both a two-state reaction model and a bivalent analyte model. Each SPR curve was successfully fitted with these models individually (local fit), yielding kinetic parameters that reflect a complex interaction. However, global fitting to these interaction models resulted in poor fits, with significant deviations observed during both the dissociation phase and towards the end of the association phase. Despite this, both the bivalent analyte and two-state conformation models estimated an apparent KD value in the nanomolar range (Table 2). Notably, the interaction between full-length AbrB and AbbA exhibited slow dissociation in the sensorgrams, indicating a high-affinity and highly stable interaction.

**Table 2.**
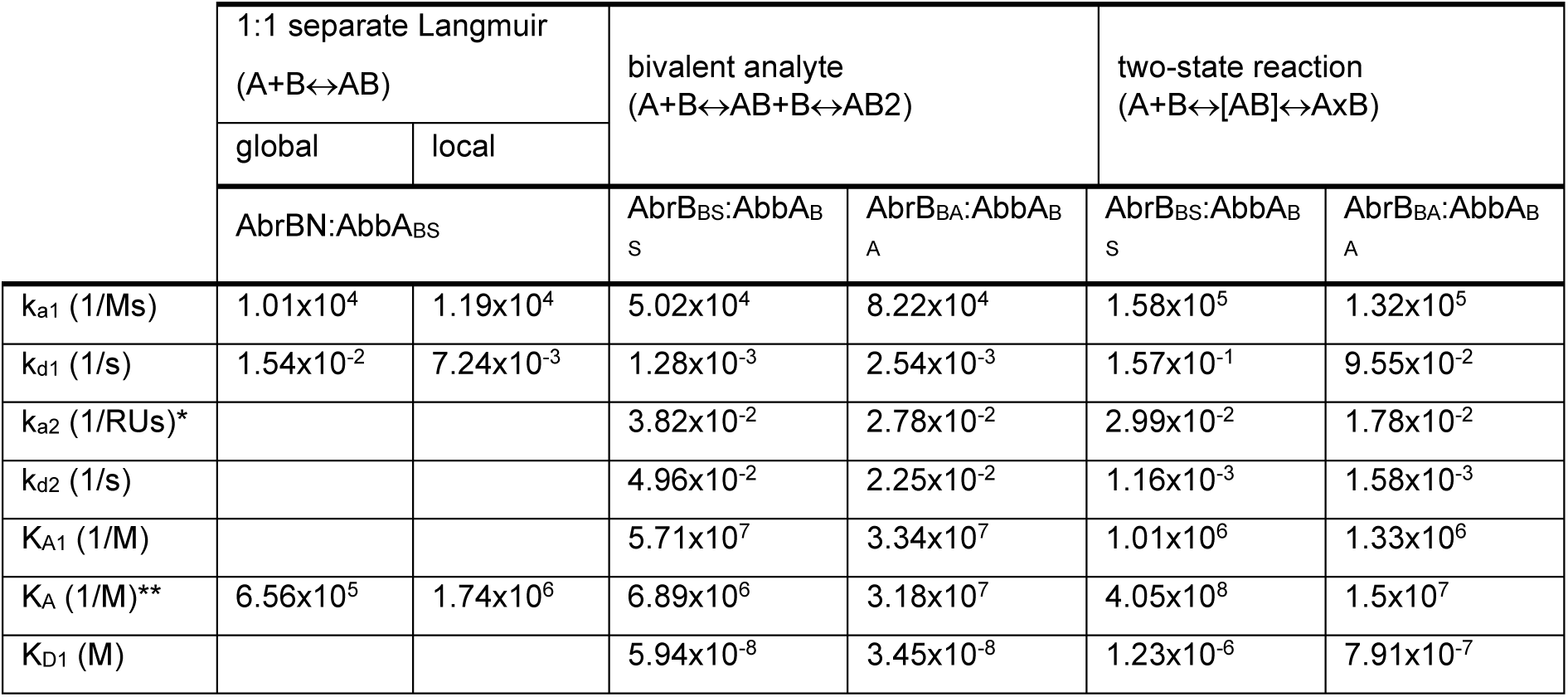
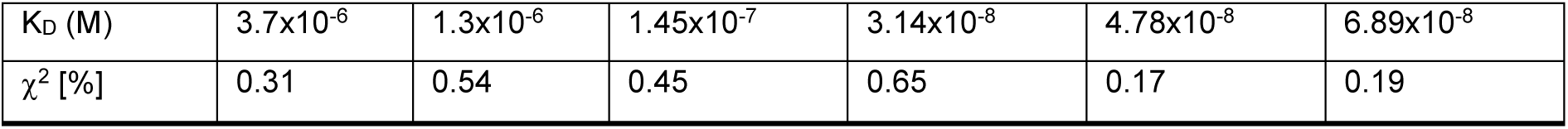
Binding parameters for the interaction between AbrB and AbbA derivates from *B. subtilis* 168 (BS) and *B. amyloliquefaciens* FZB45 (BA) as determined by surface plasmon resonance and evaluated by BIAevaluation 3.1 software using the fitting models. The truncated AbrB protein contains the N-terminal DNA-binding domain (AbrBN) from *B. subtilis* 168. The two-state reaction (conformation change) supplied directly four kinetic parameters und the apparent affinity constant K_A_ of overall reaction with the unit [M^-1^]. K_D_ was deviated from K_A_: K_D_ = 1/K_A_. The k_a2_ is independent of the molarity with the unit [s^-1^]*. The bivalent analyte model exhibits four kinetic parameters: k_a2_ present with unit [RU^-1^ s^-1^]* and KA with [RU^-1^ M^-1^]*. K_D_ and K_A_ present the equilibrium constants of overall reaction and were evaluated from the quotient of all four rate constants. K_D1_ and K_A1_ present the affinity of first reaction.

The two-state reaction (conformation change) model A + B ⇌ [AB] ⇌ A × B assumes that one molecule of AbbA binds to one AbrB tetramer, which is followed by a conformational change in the complex ([AB]), resulting in a more stable AbrB-AbbA interaction (A × B). This model produces two sets of kinetic constants: i) The association (k_a1_) and dissociation (k_d1_) rate constants describe the kinetics of the initial binding reaction; ii) The rate constants for the conformational change (k_a2_ and k_d2_) describe the subsequent stabilization of the complex.

Notably, the forward rate constant k_a2_ is independent of the molarity and is expressed in units of s^-1^. This suggests that the rate of the conformational change is consistent regardless of AbbA concentration. The kinetic parameters derived from this two-state reaction model indicate that the conformational change (second reaction) significantly stabilizes the complex formation between full-length AbrB and AbbA.

Here, the first set of rate constants reveals that the initial binding reaction is rapid but relatively unstable, with association rate constants (ka1) of 1.32×10^5^ M^-1^ s^-1^ and 1.58×10^5^ M^-1^ s^-1^, and dissociation rate constants (kd1) of 9.55×10^-2^ s^-1^ and 1.57×10^-1^ s^-1^.

Conversely, the second reaction, representing the conformational change, demonstrates a slower and more stable complex formation. The equilibrium dissociation constants (K_D_) for the overall reaction are notably lower, at 44.78×10^-8^ M for AbrB_BS_ and 6.89×10^-8^ M for AbrB_BA_, compared to the first reaction (K_D1_ of 1.23×10^-6^ M for AbrB_BS_ and 7.91×10^-7^ M for AbrB_BA_). This indicates that the conformational change stabilizes the complex, shifting the equilibrium in the forward direction.

The bivalent analyte model A + B ⇌ AB + B ⇌ AB₂ posits that each AbbA molecule can interact bivalently with one or two AbrB molecules. The first binding event is characterized by a single set of rate constants, treating the two sites on the analyte as equivalent. The second binding event, involving either a second ligand molecule or a second site on the same ligand, is described by a second set of rate constants (k_a2_ and k_d2_), which accounts for cooperative effects. These cooperative effects help explain the observed slow dissociation of the binding curves (SM 2), consistent with previous literature [43]. The association rate constant for the second site interaction is given in response units per second (RU^-1^ s^-1^) rather than in M^-1^ s^-1^, reflecting the interaction of the complex (in RU) with the ligand (in RU).

Fitting the k_a_ and k_d_ parameters can be conducted using either local or global methods, as shown in Table 2, with the 1:1 (Langmuir) model. Local fitting of the binding curves for AbbA with full-length AbrB (BS and BA) using the 1:1 model (SM 3) yielded a low χ^2^ value (χ^2^<1 % of RU_max_), indicating a good fit. However, separate fits conducted over different regions of the association and dissociation phases revealed inconsistencies in the calculated constants throughout the sensorgram. This underscores the complexity of the interaction between AbbA and full-length AbrB.

Notably, there are more pronounced differences in the KD values for AbrB_BA_ binding to AbbA_BA_ when compared using the 1:1 Langmuir model versus the complex binding model. The 1:1 Langmuir model yielded K_D_ values of 1.14×10^-6^ M or 1.24×10^-6^ M, while the complex binding model provided K_D_ values of 3.14×10^-8^ M or 6.89×10^-8^ M. These discrepancies highlight that the interaction between AbbA and AbrB_BA_ is more complex than can be described by a simple 1:1 binding model.

The separate fits of the binding curves for AbrBN with AbbA_BS_ yielded consistent rate constants for association (ka) and equilibrium constants (K_D_ and K_A_) throughout the sensorgram. However, when evaluating the AbrBN-AbbA_BS_ sensorgrams using both the two-state reaction and bivalent analyte models, we observed either a failure to fit or poor fits, as evidenced by higher χ^2^ values. These results suggest that the interaction between AbrBN and AbbA_BS_ is more straightforward compared to full-length AbrB, likely due to AbrBN’s simpler structure. AbrBN contains only one DNA-binding site [41] and interacts with a single DNA oligonucleotide without distinguishing between the GI and GII types [15]. This simplicity in the interaction is reflected in the more consistent and less complex binding kinetics.

### Parameters of negative cooperativity

In addition to kinetic analysis, we obtained thermodynamic data under equilibrium binding conditions to further characterize the interactions. Beyond the equilibrium dissociation constants, additional insights were derived from steady-state data. We plotted response units at steady-state against analyte concentration and fitted the resulting data using a general nonlinear curve fitting procedure as described previously [15]. Both the steady-state affinity model and the 4-parameter equation provided satisfactory fits with the lowest χ^2^ values .

For the interaction between AbbABS and AbrBN, the binding curve showed a hyperbolic shape (Figure 6A), consistent with a simple binding [44]. In contrast, the interaction between full-length AbrB and AbbA exhibited a curve with significant deviations from a hyperbolic shape, displaying a plane slope. This indicates a more complex interaction process between these proteins, suggesting additional factors or multiple binding events affecting the overall binding kinetics.

**Figure 6.**
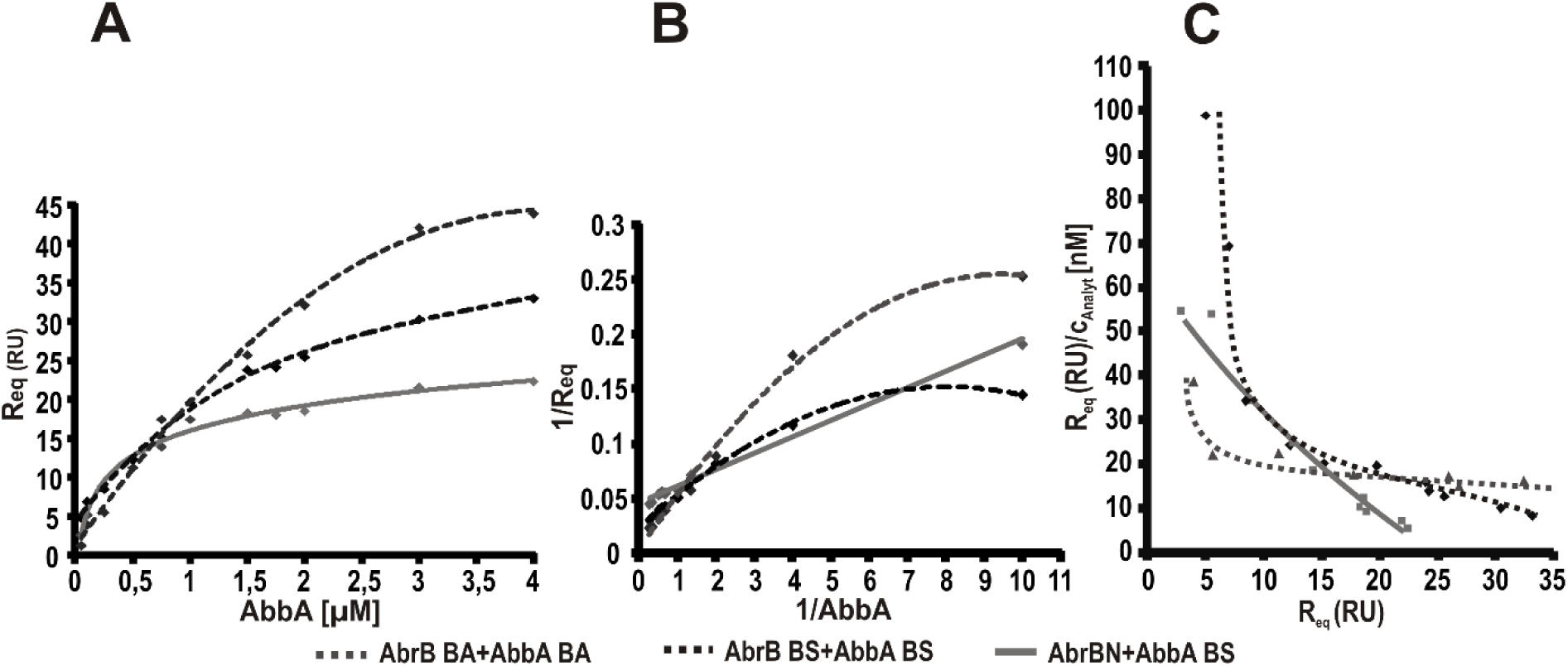
Binding isotherms of AbbA bound to AbrB, Response units at the steady-state (R_eq_) were plotted against the AbbA analyte concentration (A). Lineweaver-Burk plot (B) and Scatchard plot (C) of the isotherms illustrating the cooperative binding character (curves) of the full-length-AbrB proteins to the corresponding AbbA in contrast to the N-terminal AbrB-domains (AbrBN) (linear plots).

The presence of positive or negative cooperativity can readily be analysed from the binding isotherm using various graphical methods, including Lineweaver-Burk and Scatchard plots [45]. These plots help in diagnosing the type of cooperativity present in the binding data based on the plot’s shape (Figure 6).

The double reciprocal Lineweaver-Burk plot is used to visualize the binding data by plotting 1/R_eq_ versus 1/[AbbA]. For AbrBN interacting with AbbA, the plot is linear (Figure 6 B), which suggests simple 1:1 binding with no cooperativity. For AbrBN interacting with AbbA, the plot is linear (Figure 6 B), which suggests simple 1:1 binding with no cooperativity.

The binding curve of full-length AbrB at AbbA indicated a common shape for negative cooperativity. The Scatchard plot was constructed by plotting R_eq_/[AbbA] [versus R_eq_, where R_eq_ is the equilibrium response units (RU) and [AbbA] is the analyte concentration [nM]. For AbrBN-AbbA_BS_, the Scatchard plot is almost a straight line, indicating no cooperativity. In contrast, the Scatchard plot for the interaction of full-length AbrB_BS_ with AbbA_BS_ displays a curvilinear (concave) shape (Figure 6 C), which is characteristic of negative cooperativity [45–47]. This suggests that the binding affinity changes as the reaction progresses, showing that the interaction is not uniform across the saturation range.

These analyses confirm that while AbrBN exhibits simple binding without cooperativity, the interaction between full-length AbrB and AbbA involves negative cooperativity, affecting how the affinity of the binding changes with the degree of saturation.

To further characterize the cooperativity in binding interactions, we utilized Hill plots, where log[R_eq_/(RU_max_-R_eq_)] as a function of analyte concentration log[AbbA]. The RU_max_ values used in these plots were derived from either the steady-state affinity (ss) or 4-parameter equation (pe) fitting models, providing two Hill coefficients (n_H_) for each AbrB-AbbA interaction pair.

In contrast, for the binding of AbbA_BS_ to AbrB_BS_, we observed Hill coefficients of n_H_^(ss)^=0.77 and n^H(pe)^=0.89, and for AbbA_BA_ binding to AbrB_BA_, the Hill coefficients were n_H_^(ss)^=0.93 and n_H_^(pe)^=0.75. These coefficients, particularly those derived from the 4-parameter equation fitting model, were distinctly below 1 (n_H_<1), indicating negative cooperativity in these binding systems. Negative cooperativity suggests that while the binding sites are identical, interactions between them result in a scenario where binding at one site decreases the affinity of the remaining sites [46]. This type of cooperative behavior implies a more complex interaction dynamic, where the binding of one AbbA molecule to AbrB influences the binding affinity of subsequent AbbA molecules.

### Analysis of the conformational changes of proteins

Circular dichroism (CD) spectroscopy is a powerful tool for detecting protein conformational changes induced by DNA binding and interactions with other binding partners. Figure 7 presents the CD spectra of the AbrB_BS_ protein in its free form, as well as in complexes with DNA and AbbA. The measurements were conducted across a wavelength range of 260 to 185 nm, focusing on the far-UV spectrum to monitor the secondary structure of the protein. Due to significant spectral noise below 205 nm - attributable to the use of a 1 mm quartz cuvette necessary for large sample volumes, as well as the high protein concentrations required for these binding studies - the data from this region were excluded from the analysis. Consequently, CD spectroscopy was utilized in this study for qualitative structural characterization rather than precise quantification of secondary structure elements.

**Figure 7.**
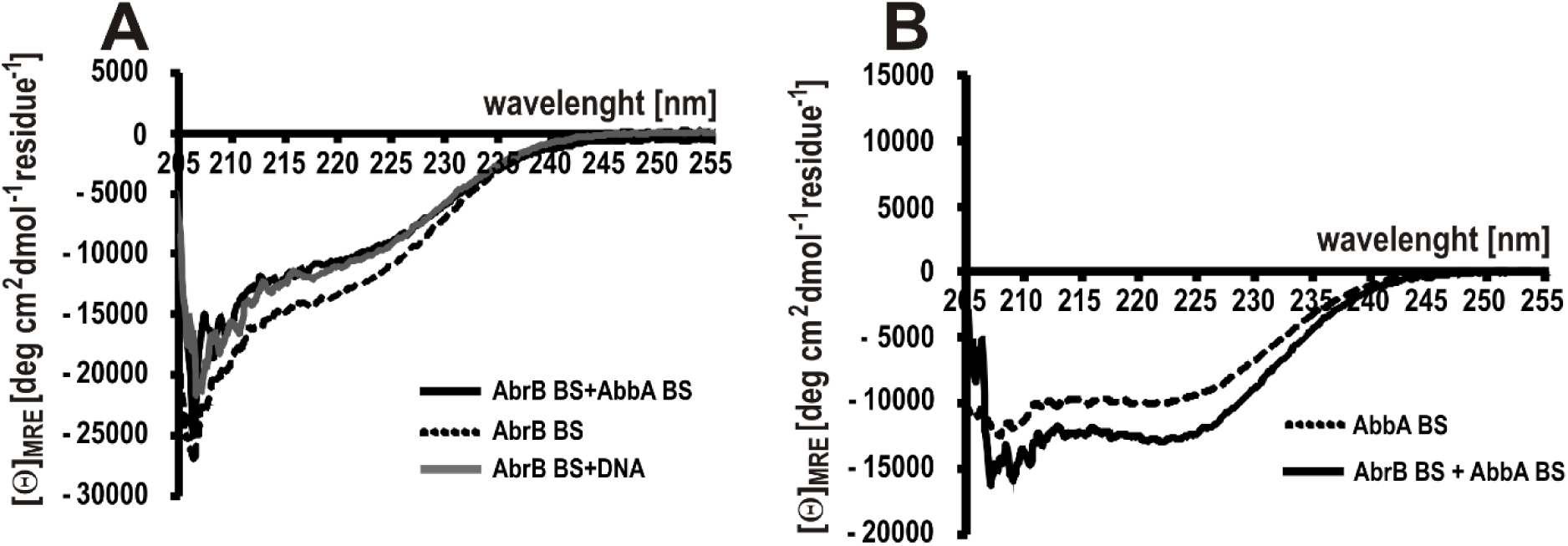
Near- and far-UV CD-spectra of the proteins, DNA-AbrB_BS_ and AbrB_BS_-AbbA_BS_ complexes. All measurements were performed in triplicate at room temperature. (A) Comparison of the free AbrB protein (dashed line) and the AbrB protein in the AbrB_BS_-DNA complex (grey line) or AbrB_BS_-AbbA_BS_ complex (black line). Spectra of free DNA or AbbA_BS_ were subtracted respectively resulting in spectral contributions of the bound AbrB_BS_. (B) Comparison of free AbbA_BS_ protein (dashed line) and bound to AbrB_BS_ (black line) (subtracted spectrum of AbbA_BS’_).

While the complexity of the AbrB_BS_ CD data prevented a detailed deconvolution into specific secondary structures via the DichroWeb server, reference spectra from previous studies [25, 48, 49] suggest that the AbrB_BS_ protein exhibits a mixture of both α-helical and β-sheet structures. The CD spectra for AbbA_BS_, on the other hand, are characteristic of proteins with predominantly α-helical content. These observations align with known structural features of these proteins, further supporting the integrity of the experimental approach and the qualitative conclusions drawn from the CD data.

In the binding studies, AbrB_BS_ protein (20 µM) was analyzed in complex either with a group I ds-DNA oligonucleotide named D4 (5 µM) or with its antirepressor AbbA_BS_, which was present in excess at a concentration of 40 µM. The group I ds-oligonucleotides, recognized as DNA targets of AbrB, were previously shown to bind AbrB with higher affinity and cooperativity, as confirmed by SPR binding kinetics [15, 23]. Given that AbrB forms tetramers and AbbA exists as a dimer (as evidenced by FPLC data in Figure 1), the concentrations used in this study corresponded to a 1:4 molar ratio of AbrB4 to AbbA2, which indicates an excess of AbbA (1 AbrB4: 4 AbbA2). This ratio was determined to be necessary by gel shift assays (Figure 5), which showed that at least a fourfold concentration of AbbA dimers was required to dissociate the AbrB4-DNA complex.

The CD spectra revealed that the interaction of AbrB with either the antirepressor AbbA or the D4 oligonucleotide induced similar conformational changes in the AbrB protein (Figure 7A). Both AbbA and DNA caused comparable alterations in the ellipticity of the AbrB spectrum, particularly between 205 and 230 nm, which suggests that the binding of these molecules to AbrB influences its secondary structure in a similar manner.

Additionally, the presence of AbrB induced conformational changes in the AbbA protein as well (Figure 7B). The typical α-helical spectrum of AbbA was altered, leading to an increased negative ellipticity near 222 nm and 208 nm. This suggests that the interaction with AbrB not only stabilizes the complex but also affects the secondary structure of AbbA, further supporting the idea of a dynamic interaction between these two proteins.

### Protein secondary structure analysis

To deconvolute the protein CD data into secondary structure elements of AbrB and AbbA proteins, we utilized the online DichroWeb server [25]. This server allows for the analysis of CD spectra using five widely recognized algorithms. For our analysis, we employed three specific algorithms: SELCON3 [26–28], CONTIN [29, 30], and CDSSTR [28, 31] Each of these programs requires the use of reference datasets, which are constructed from the spectra of proteins with known crystal structures and, consequently, well-characterized secondary structures [48].

Given that the CD signal becomes increasingly noisy at shorter wavelengths, we limited our analysis to wavelengths above 185 nm to ensure data reliability. The analysis was performed on both AbrB and AbbA proteins using reference datasets 3, 4, 6, 7, and SP175, which are available on the DichroWeb platform. A key parameter in this analysis is the normalized root mean square deviation (NRMSD), which measures the fit between the experimental and calculated spectra. The NRMSD serves as an indicator of the quality of the deconvolution results. In our study, we accepted the calculated secondary structures only when the NRMSD values were below established thresholds, specifically NRMSD < 0.1 for the CDSSTR and CONTIN algorithms, and NRMSD < 0.258 for the SELCON3 algorithm. These criteria ensured that the deconvoluted secondary structures were as accurate as possible, reflecting a good fit between the observed CD data and the predicted models.

The CD measurement of both AbrB proteins of *B. subtilis* and *B. amyloliquefaciens* FZB45 revealed identical spectra, characterized by two unequal minima at approximately 206 nm and 219 nm (Figure 8A), consistent with previously reported data for AbrB_BS_ [50]. However, the CD spectrum of the truncated AbrBN protein exhibited a shift in the minimum towards the far-UV wavelength region (∼203 nm) and presented a shoulder-like feature around 214 nm.

**Figure 8.**
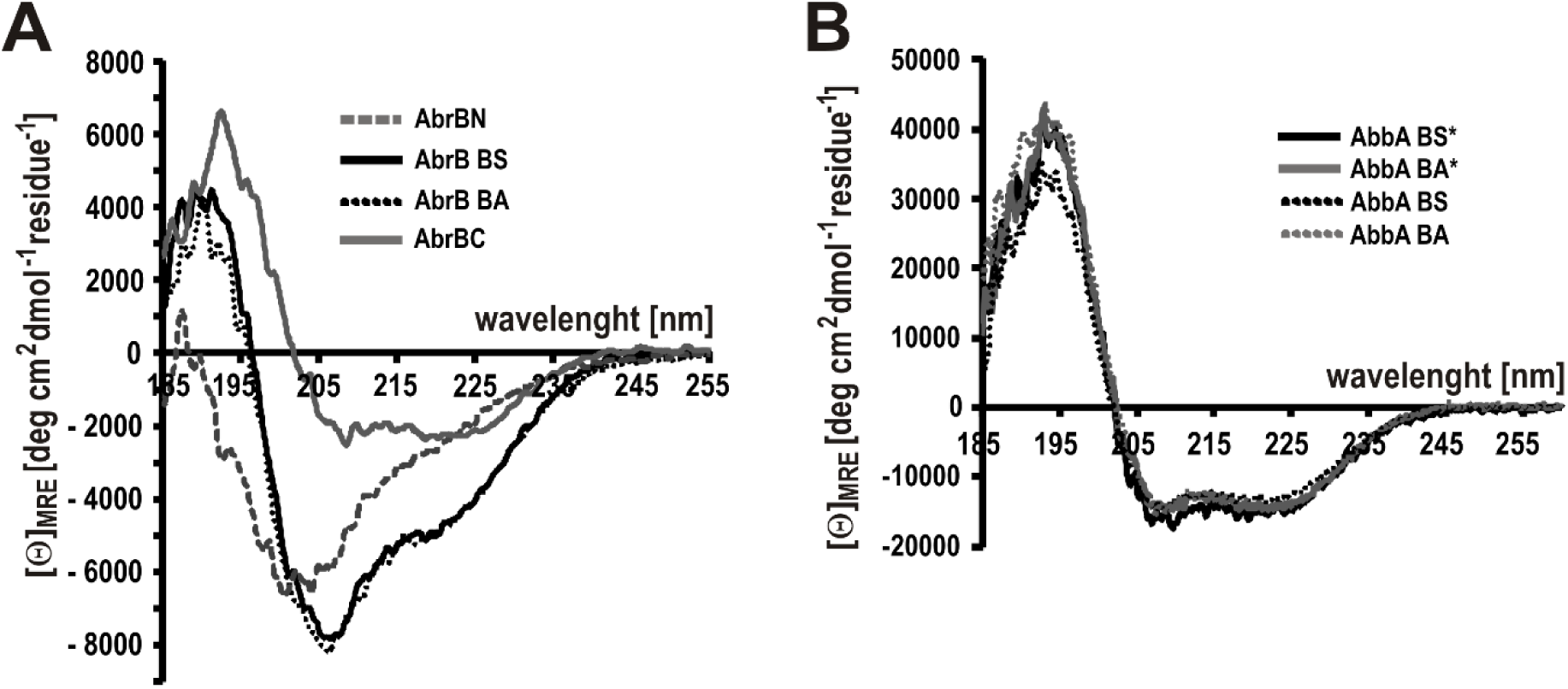
Near- and far-UV CD spectra of AbrB (A) and AbbA (B) variants. Twenty scans for each protein were accumulated and averaged. AbrBN represents the N-terminal DNA-binding domain of AbrB protein. The AbrBC spectrum was obtained as a difference spectrum of the full-length AbrB_BS_ and AbrBN. * His-tagged AbbA-variants exhibited nearly identical CD-spectra compared to the variants with truncated His-tag indicating that the His-tag does not significantly impact on secondary structure.

The CD spectra for both full-length AbrB proteins demonstrated a complex structural composition, with a mixture of β-sheet elements indicated by a minimum at 219 nm and α-helices identified by a distinct maximum around 190 nm, along with a negative peak near 209 nm. The spectra, particularly the low signal intensity at 209 nm for AbrB and 203 nm for AbrBN, suggest an atypical α-helical folding. This is indicative of a low α-helix content, and instead, it points towards a significant presence of β-turns and natively disordered structures [51]. This conclusion is supported by the analysis of the secondary structure elements, which are detailed in Table 3.

**Table 3:** Deconvolution of protein CD data into secondary structure elements of AbrB and AbbA proteins using algorithms CDSSTR, CONTIN and SELCON3 as well as the use of different reference datasets 3, 4, 6, 7 and SP175. The protein total length is 114 amino acids for full-length AbrB, 75 aa for AbrBN including the His_6_-tag and 39 aa for C-terminal domain (AbrBC). The content of disordered structures bracketed refers to AbrB proteins lacking the 20 aa long His_6_-tag. BS = *B. subtilis* 168 and BA = *B. amyloliquefaciens* FZB45. *The percentage for secondary structure of AbrBN is derived from NMR structure [41]. ^#^The percentage for secondary structure of AbrBC including 42 aa is derived from NMR structure [53].

| | | α-helix | | β-sheet | | β-turn | | natively disordered | |
| --- | --- | --- | --- | --- | --- | --- | --- | --- | --- |
|  |  | AS | % | AS | % | AS | % | AS | % |
| AbrBN* |  | 7 | 13.2 | 25 | 47.2 | 12 | 26.6 | 9 | 17 |
| AbrBN |  | 4.4 | 5.8 | 25 | 33.5 | 14 | 18.3 | ~31<br>(11) | ~42<br>(15) |
| AbrB<sub>BS</sub> |  | 16.2 | 14.2 | 33.3 | 29.2 | 21.1 | 18.5 | ~42<br>(22) | ~37<br>(19) |
| AbrB<sub>BA</sub> |  | 12 | 10.5 | 37.5 | 32.9 | 22.2 | 19.5 | ~42<br>(22) | 37 (19) |
| AbrBC |  | 12 | 30.8 | 8 | 22.2 | 7.5 | 19.2 | 11 | 28.2 |
| AbrBC<sup>#</sup> |  | 15 | 35.7 | 6 | 14.3 | 6 | 14.3 | 15 | 35.7 |
| AbbA<sub>BS</sub> |  | 40.3 | 59.3 | 6.5 | 9.6 | 8.1 | 11.9 | 13.1 | 19.3 |
| AbbA<sub>BA</sub> |  | 38.5 | 56.6 | 7.5 | 11 | 8.8 | 12.9 | 13.5 | 19.8 |

The analysis showed that all three AbrB proteins - AbrBN, AbrBBS, and AbrBBA - exhibited a high β-sheet content, ranging from 29% to 33.5%. Full-length AbrBs displayed a higher α-helical content compared to the truncated AbrBN. It is important to note that these CD measurements were performed on AbrB variants that included a 20 amino acid long His6-tag fused at the N-terminus. The His6-tag, which constitutes 26.5% of AbrBN and 17.5% of the full-length AbrB proteins relative to their total length, is known to form a disordered, extended structure, as described in various recombinant proteins [52].

When comparing the truncated AbrBN from this study to AbrBN_53_, whose NMR structure was refined by Sullivan and colleagues [41], only slight differences were observed in the distribution of disordered structures (9 to 11 amino acids). Both AbrBN proteins exhibited similar amounts of amino acids involved in β-sheets (25 residues), β-turns (12 to 14 residues), and α-helices (4 to 7 residues). The full-length AbrB proteins also showed nearly identical distributions of amino acids in β-turns, disordered structures, and well-defined secondary structures, underscoring the consistency of structural elements across these variants.

In addition to analyzing the full-length AbrB_BS_ and the truncated AbrBN proteins, we derived the secondary structures specifically for the C-terminal domain by deconvoluting the CD spectra of these proteins. The deconvolution results allowed us to determine the amino acid content and the percentage of each secondary structure element present in the C-terminal domain (Table 3). The CD studies of this domain revealed a mixture of well-defined secondary structures, including β-sheets, β-turns, and α-helices, alongside disordered regions.

To further characterize the C-terminal domain, we generated a CD spectrum by subtracting the spectrum of AbrBN from that of the full-length AbrB_BS_. This difference spectrum exhibited features indicative of helical structures, although the low ellipticity observed at the characteristic minima (near 222 nm and 208 nm) and at the maximum (around 192 nm) suggested the presence of other secondary structures in addition to α-helices.

Interestingly, this observation aligns with findings from a study by Tucker A.T., who reported a similar CD spectrum and ellipticity distribution for the cloned and purified AbrBC from *B. subtilis* [20]. Moreover, the NMR structure of AbrBC, as described by Olson in 2013 [53], which comprises 42 amino acid residues, also indicated a similar distribution of secondary structures, further supporting our CD analysis of the C-terminal domain (Table 3). These results highlight the structural complexity of the C-terminal domain, emphasizing the presence of multiple secondary structure elements beyond just α-helices.

Our own preliminary study [54] provided the first CD spectra of AbbA from *B. subtilis*, revealing its canonical alpha-helix secondary structure. Building on this, we compared the CD spectra of AbbA proteins from *both B. subtilis* and *B. amyloliquefaciens* FZB45. The CD measurements for both AbbA proteins yielded very similar spectra and ellipticity distributions, which was expected given their high amino acid identity of approximately 89%.

The spectra of both AbbA proteins are consistent with a predominantly α-helical structure, characterized by two negative ellipticity peaks at around 208 nm and 219 nm, and a positive ellipticity maximum at 192 nm (Figure 8 B). These spectral features align with the intense CD signals typically produced by helical components, as described previously [48]. The secondary structure analysis revealed that both AbbA proteins, which lack His6-tags, contain a high helical content, approximately 60%, corresponding to 38 to 40 amino acids of the total protein length (Table 3).

While the His6-tagged AbbA proteins exhibited discrepancies in the deconvolution of their secondary structures (data not shown), previous in vitro binding experiments demonstrated that the presence of the His6-tag did not affect the function of AbbA. This suggests that the structural differences observed in the His6-tagged versions may not impact the biological activity of AbbA, though they do alter the CD spectra.

### SAXS structure of AbrB and AbbA

Small-angle X-ray scattering (SAXS) is a well-established method for obtaining low-resolution structural information about biological macromolecules in solution. The technique generates scattering patterns from a dilute protein solution, which are typically presented as radially averaged one-dimensional curves (see SM 6). These curves provide important parameters (detailed in SM 5), such as the size, oligomeric state, and overall shape of the molecule. These parameters can be determined ab initio, offering insights into the macromolecular structure (as detailed in the Materials and Methods section).

In this study, we employed the popular ab initio program GASBOR to reconstruct low-resolution three-dimensional structures based on an assembly of dummy residues [33, 40]. GASBOR enabled us to determine the shape and fold of AbrB and AbbA proteins at a resolution of approximately 0.5 nm. However, it is important to note that solution scattering, such as SAXS, is not suitable for determining secondary structure elements. To account for the bound solvent, the proteins were modeled with a surrounding hydration layer, which made them appear larger in the SAXS data.

Figure 9 illustrates the SAXS-derived structures of the AbrB tetramer and the AbbA dimer, as reconstructed using GASBOR. Additionally, the Guinier analysis performed with the PRIMUS program provided molecular masses (kDa) of the proteins that correlate with their oligomeric forms (details in SM 5). The molecular mass of the AbbA dimer, as determined by SAXS, was 16 kDa, which closely matches the theoretical mass of 15.78 kDa. Similarly, the molecular mass of the AbrB tetramer was found to be approximately 40 kDa, which is close to the theoretical mass of 50.56 kDa. The structural models were constructed assuming P22 particle symmetry for the AbrB tetramer and P2 symmetry for the AbbA dimer, reflecting their respective oligomeric states.

**Figure 9.**
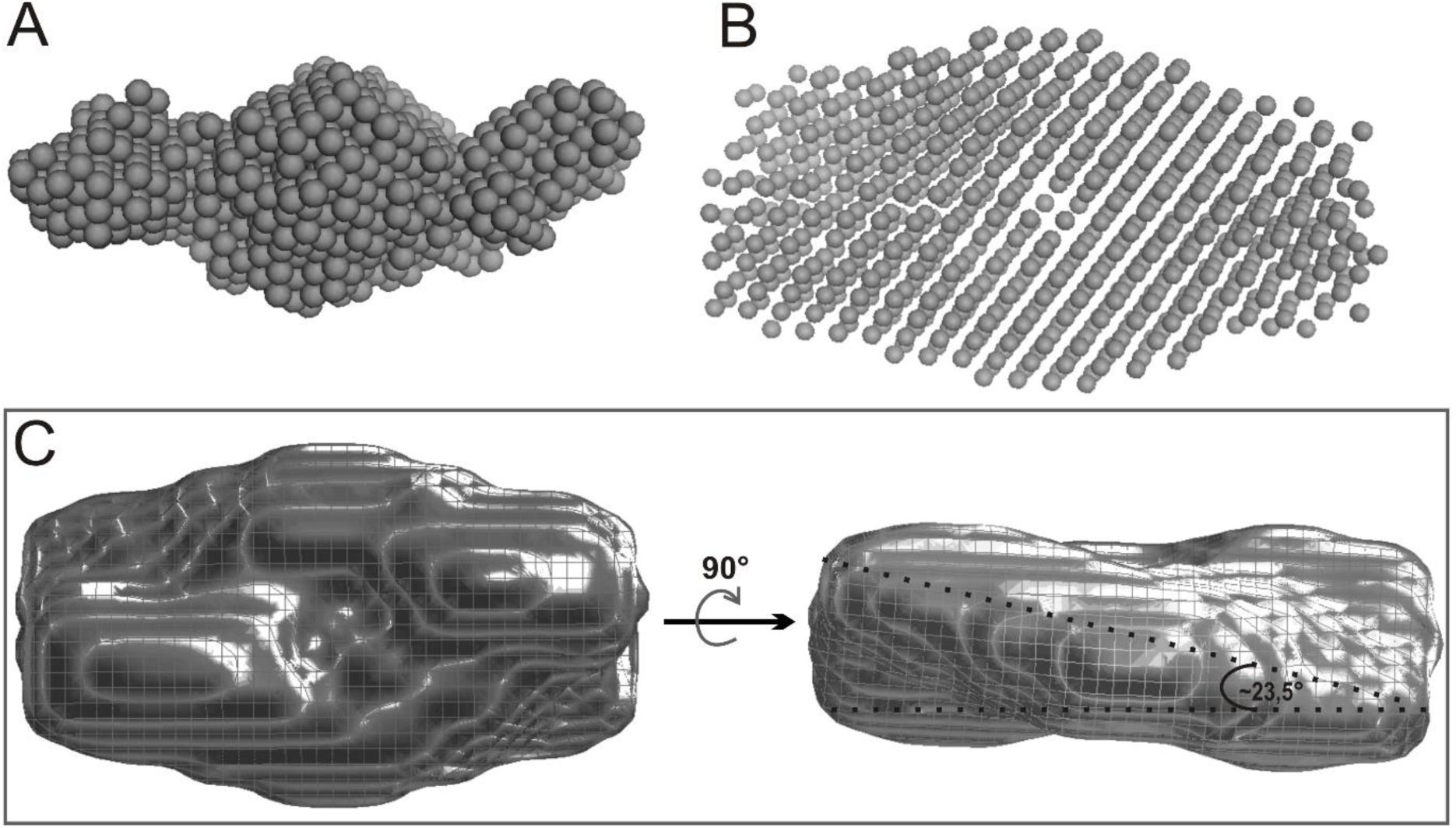
SAXS structures of AbbA dimer (A) and AbrB tetramer (B) represented as spheres model. Low resolution structures of proteins calculated from scattering data with *ab initio* GASBOR program assuming P22 symmetry for AbrB tetramer and P2 for AbbA dimer. (C) The dummy residues of AbrB tetramer are covered of mesh, surface and subsequently represented as wireframe graphics. The view rotated 90°.

The works of Olson et al. presented the first structural model of the full-length AbrB by using the NMR structure of the C-terminal domain (AbrBC) and subsequently orienting the N-terminal domain (AbrBN, PDB: 1Z0R) and AbrBC relative to each other based on distance restraints obtained from chemical crosslinking and LC/MS/MS analysis[8]. Despite this progress, the overall structure of the AbrB tetramer remains unknown. This is largely due to the highly dynamic nature of the full-length AbrB protein, which has so far resisted crystallographic and NMR-based structural determination.

Attempts to crystallize AbrB in the presence of two strongly binding 25-mer oligonucleotides were unsuccessful, likely due to the flexibility of the C-terminal domain, which hinders stable crystal formation. Given these challenges, we turned to small-angle X-ray scattering (SAXS) to investigate the overall conformation of the full-length AbrB under native conditions.

Our SAXS analysis proposes the first overall structure of the AbrB tetramer, which appears to have an oblong shape with dimensions of approximately 111 Å between the C-termini and 59 Å between the N-termini. The N-terminal DNA-binding domains are positioned symmetrically and antiparallel to each other, while the C-terminal domains are rotated by about 23.5° relative to each other. This SAXS-derived structural model is consistent with previous studies by Benson (2002), which suggested that the AbrB tetramer has a non-spherical shape, leading to an increased Stokes radius during gel filtration. This complements the modeled full-length structure of AbrB recently published by others .

In contrast to the symmetric AbrB tetramer, the antirepressor AbbA dimer exhibited a significantly different disposition. The AbbA dimer has a more elongated shape, with dimensions of approximately 66 Å by 30 Å, and shows pseudo-tandem symmetry. Interestingly, the AbbA dimer is not symmetrically arranged; instead, it is rotated about its longitudinal axis, which results in asymmetrically oriented side arms. This distinctive structural arrangement of the AbbA dimer further highlights the differences in conformational dynamics and binding interactions between AbrB and AbbA (see SM 7).

### Model of AbrB-AbbA interactions

Our previous findings support a model where multiple AbrB tetramers simultaneously bind to DNA by looping and interacting with both strands through their two DNA-binding sites at opposite ends of the tetramer [15]. This interaction involves complex, cooperative effects, especially in the *phyC* promoter region, where extensive protection by AbrB was observed in footprint assays. The C-terminal domains of AbrB are crucial for these cooperative interactions, as AbrBN alone does not exhibit cooperativity (Figure 10, first complex with AbrB).

**Figure 10.**
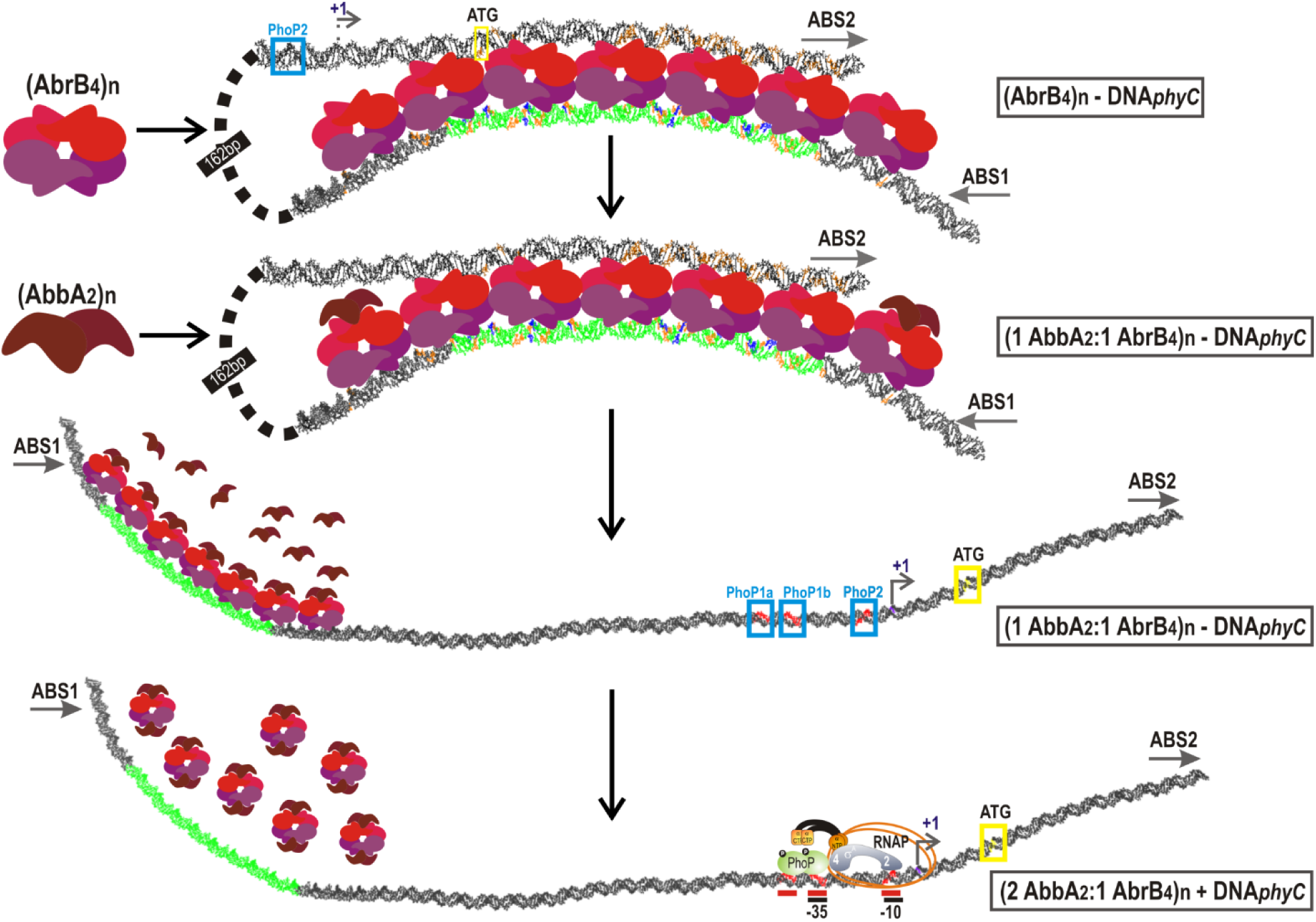
Proposed mechanism of AbrB.interactiona with phyC promoter and the antirepressor activity of AbbA.

Most likely, AbbA functions as an antirepressor by mimicking the structure of *phyC* DNA. A similar mechanisms was proposed for CarS antirepressor. According to Leon et al., the CarS mimics operator DNA to sequester the DNA-recognition helix of the CarA repressor [55]. CarS achieves this by presenting a polar surface similar in size and charge distribution to the operator DNA, preventing CarA from binding.

AbbA appears to use a similar mechanism. It is rich in negatively charged residues and has a theoretical isoelectric point (IEP) of around 4.5, which suggests it may mimic DNA structure. Furthermore, SAXS analysis indicates that the AbbA dimer resembles a deformed DNA double helix, supporting the idea that AbbA sequesters AbrB’s DNA-binding sites by mimicking DNA. Our findings also showed that AbbA does not bind to AbrB’s C-terminal domain, reinforcing the idea that AbbA competes with DNA for AbrB’s binding sites. SPR binding kinetics reveal that these domains facilitate conformational changes within the AbrB-AbbA complex. AbbA’s dimeric nature likely allows it to bind two AbrB molecules or mimic the DNA-binding site on AbrB. Our data show that AbbA exhibits negative cooperativity in its interactions with AbrB, as demonstrated by Scatchard plots and Hill coefficients. This indicates that the binding of the first AbbA dimer to AbrB occurs with high affinity and induces a conformational change that reduces the affinity for a second AbbA dimer.

AbrB, in its tetrameric state, possesses two DNA-binding sites [24, 56], with one site formed by two N-terminal domains. Therefore, we hypothesize that one AbbA dimer interacts with a single DNA-binding site of AbrB, which consists of two N-terminal halves. This idea is supported by the well-fitting Langmuir model, which describes a simple 1:1 interaction for AbrBN-AbbA interactions with a micromolar affinity of 3.7×10^-6^ M.

## Conclusions

The interaction between AbrB and AbbA plays a critical role in regulating gene expression in *Bacillus* species, specifically in the repression of the *phyC* gene. Similar to other promoters like *tycA* [57]., the binding of AbrB to DNA is cooperative, with multiple AbrB tetramers involved, suggesting a complex regulatory mechanism.

Non-covalent binding to an antirepressor like AbbA can inactivate a repressor through different mechanisms, including the occlusion of DNA-binding elements. SPR experiments have shown that AbrB binds strongly to DNA, with minimal dissociation over time, raising the question of how AbrB dissociates from DNA *in vivo*. The antirepressor AbbA has been identified as a key player in this process, blocking AbrB from binding to target gene.

Our *in vitro* studies confirm that AbbA acts as a DNA competitor, preventing AbrB from binding to the *phyC* promoter in a concentration-dependent manner. This was evident in gel shift experiments where AbbA reversed AbrB’s binding to DNA. Notably, AbbA does not bind directly to the *phyC* promoter region, but rather interacts with AbrB to disrupt its DNA-binding ability. CD and SPR techniques revealed that DNA and AbbA compete for the same binding site within the AbrB tetramer, a finding supported by the loss of binding in an AbrB mutant where the conserved arginine residue (R15) was substituted.

Recent studies suggest that AbbA may function similarly to the bacterial antirepressor CarS, which mimics operator DNA to sequester the DNA-recognition helix of the CarA repressor. AbbA, with its high content of negatively charged residues, could mimic the structure of the *phyC* DNA, facilitating its interaction with AbrB. This idea is supported by SAXS data showing that the overall shape of the AbbA dimer resembles a deformed DNA double helix, consistent with the hypothesis that AbrB recognizes specific DNA structures rather than sequences.

ITC and SPR data have shown that AbbA does not bind to the C-terminal domain of AbrB (AbrBC), instead displacing AbrB from DNA by occupying its DNA-binding sites. Differences in the dissociation kinetics of AbrB from the *phyC* region in the presence of AbbA suggest nuanced interactions, potentially influenced by variations in the AbbA proteins from different *Bacillus* strains.

Further SPR studies revealed that the interaction between full-length AbrB and AbbA is more complex than previously understood. A two-state reaction model and a bivalent analyte model described the interaction, with nanomolar affinities for AbrB and AbbA from different *Bacillus* species. Interestingly, the interaction exhibited negative cooperativity, where the apparent affinity of AbrB for AbbA decreased as the binding sites became saturated. This is reminiscent of the negative cooperativity observed in insulin receptor interactions with insulin.

Scatchard plots of the AbrB-AbbA interaction showed curvilinearity, indicating the presence of both high- and low-affinity binding sites within the AbrB tetramer. The binding of the first AbbA dimer to AbrB likely induces a conformational change, reducing the affinity for the second AbbA dimer. This effect was particularly pronounced in the AbrB-AbbA interaction from B*. amyloliquefaciens* FZB45, where the dissociation of the bound analyte from the ligand accelerated - a hallmark of negative cooperativity.

Our SAXS analysis of the AbrB tetramer, combined with previous structural models, supports the idea that AbrB possesses symmetric, antiparallel DNA-binding sites. This arrangement may explain the observed negative cooperativity and the complex kinetics of the AbrB-AbbA interaction. The antiparallel symmetry and cooperative binding of AbrB suggest a sophisticated regulatory mechanism that ensures precise control over gene expression during the transition state in *Bacillus* species.

Finaly, we cnclude that AbbA mimics DNA to displace AbrB from its DNA-binding sites, leading to the activation of transition-state genes. SAXS and structural data support this mechanism, revealing that AbbA and AbrB interact in a manner analogous to DNA-protein interactions, with implications for understanding transcription regulation in bacteria.

## Acknowledgements

This work was supported by Deutsche Forschungsgemeinschaft, DFG (grant BO 1113/9-1 to R.B.). Our special thanks go to Sabine Nicklisch and Christiane Müller for skillful technical assistance. The authors thank Tobias Werther and Heike Nikolenko from FMP (Leibniz-Institut für Molekulare Pharmakologie, Berlin, Germany) for access to circular dichroism spectrometer and technical support. SPR measurements were performed in laboratory of FMP with support from Tobias Werther. SAXS measurements and evaluation were carried out in EMBL (DESY, Hamburg, Germany) with support from Manfred Roessle. The authors also thank Olga Dolgova and Maria Tsyglakowa for support of construction of plasmids and overexpression of proteins.

## Abbreviations

DTT: 1,4-dithio-D-threitol
BCIP: 5-bromo-4-chloro-3-indolyl phosphate
NBT: nitro blue tetrazolium chloride
DMF: dimethylformamide
SDS: sodium dodecyl sulfate
SPR: surface plasmon resonance
ABS: AbrB-binding site

## Footnotes

1 http://smart.embl-heidelberg.de/smart/show_info.pl

