## Supplementary Material for "AbrB antirepressor AbbA is a competitive inhibitor of AbrB-*phyC* interaction"

Mailing address:

Oliwia Makarewicz

Universitätsklinikum Jena

Institute of Infectious Diseases and Infection Control

Am Klinikum 1

D-07747 Jena, Germany.

**SM1.** Gel shift assays with increasing AbbA concentrations without AbrB in presence of the 523 bp *phyC* DNA region (-392 to +131, Sn4/ F3rev-PCR product, 10000 cpm corresponding to 0.37 nM) was used for control assay. The control lanes (0 µM) are without AbbA protein. White arrowheads, free DNA.

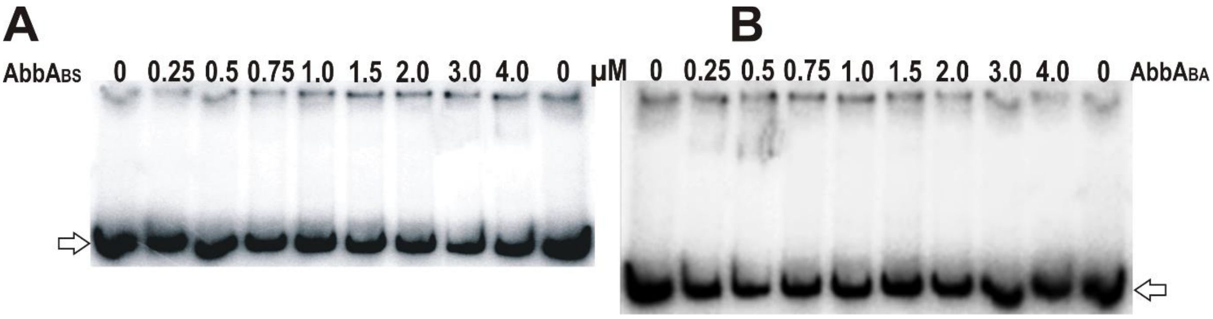

**SM2.** SPR sensorgrams of AbrB and AbbA interactions. The binding of soluble AbbA from *B. subtilis* (BS) and *B. amyloliquefaciens* FZB45 (BA) to immobilized AbrBN (A) as well as the full-length AbrB$_{BS}$ (B) and AbrB$_{BA}$ (C) was examined using a Biacore 3000 optical biosensor equipment (Biacore, GE Healthcare). AbrB proteins were covalently immobilized to the carboxyl methylated dextran surface of a CM5 sensor chip (AbrB$_{BA}$ at the flow cell Fc4; AbrB$_{BS}$ at Fc3; AbrBN at Fc2), and AbbA protein at the indicated concentrations was injected into the flow-cells, with binding being monitored for 135 seconds, followed by a 115-seconds dissociation period. We used a flow-cell within the CM5 sensor chip with no immobilized AbrB as a control cell (Fc1).

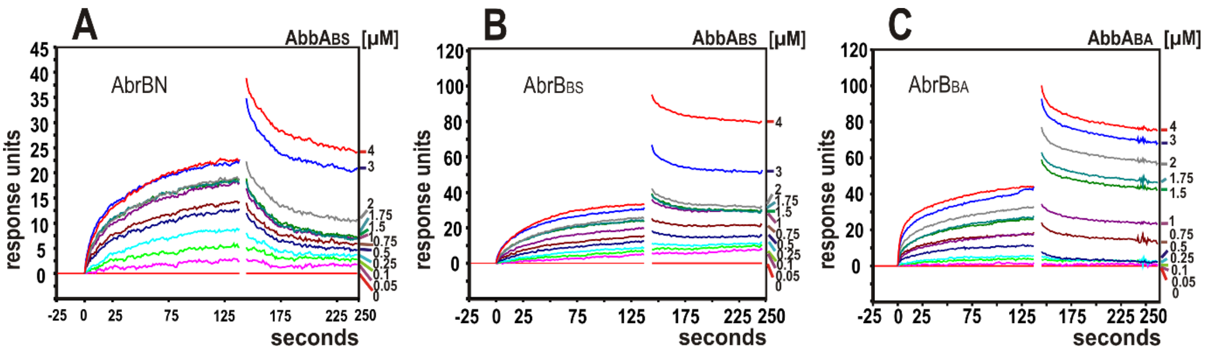

**SM3.** Binding curves and parameters (table below) for the interaction between AbrB and AbbA proteins from *B. subtilis* 168 (BS) and *B. amyloliquefaciens* FZB45 (BA) as determined by SPR and evaluated by BIAevaluation 3.1 software using the fitting models: global or local separate fitting 1:1 (Langmuir) model, two-state reaction (conformation change) model (A+B ↔ [AB] ↔ A×B) and bivalent analyte model (A+B ↔ AB+B ↔ AB2). The results of fitting were accepted when $\chi^2$<1 % of $RU_{max}$.

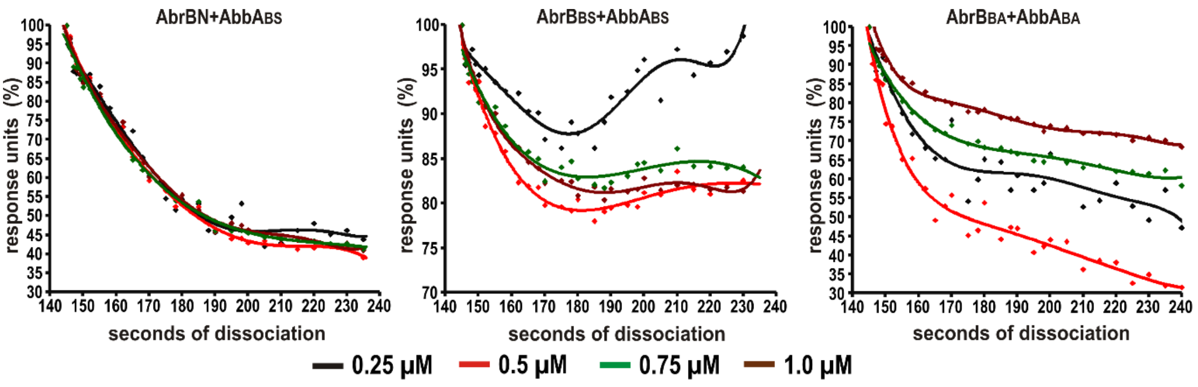


| | | 1:1 separate Langmuir (A+B↔AB) | | bivalent analyte (A+B↔AB+B↔AB2) | two-state reaction (A+B↔[AB]↔AxB) |
|---|---|---|---|---|---|
| | | global | local | | |
| $AbrB_{BS}$:$AbbA_{BS}$ | $k_{a1}$ (1/Ms) | $2.49 \times 10^{4}$ | $1.99 \times 10^{4}$ | $5.02 \times 10^{4}$ | $1.58 \times 10^{5}$ |
| | $k_{d1}$ (1/s) | $7.2 \times 10^{-3}$ | $3.09 \times 10^{-3}$ | $1.28 \times 10^{-3}$ | $1.57 \times 10^{-1}$ |
| | $k_{a2}$ (1/RUs)* | | | $3.82 \times 10^{-2}$ | $2.99 \times 10^{-2}$ |
| | $k_{d2}$ (1/s) | | | $4.96 \times 10^{-2}$ | $1.16 \times 10^{-3}$ |
| | $K_{A1}$ (1/M) | | | $5.71 \times 10^{7}$ | $1.01 \times 10^{6}$ |
| | $K_A$ (1/M)** | $3{,}72 \times 10^{6}$ | $8.55 \times 10^{6}$ | $6.89 \times 10^{6}$ | $4.05 \times 10^{8}$ |
| | $K_{D1}$ (M) | | | $5.94 \times 10^{-8}$ | $1.23 \times 10^{-6}$ |
| | $K_D$ (M) | $7.85 \times 10^{-7}$ | $2.96 \times 10^{-7}$ | $1.45 \times 10^{-7}$ | $4.78 \times 10^{-8}$ |
| | $\chi^2$[%] | 0.39 | 0.88 | 0.45 | 0.17 |
| AbrBN:$AbbA_{BS}$ | $k_{a1}$ (1/Ms) | $1.01 \times 10^{4}$ | $1.19 \times 10^{4}$ | | |
| | $k_{d1}$ (1/s) | $1.54 \times 10^{-2}$ | $7.24 \times 10^{-3}$ | | |
| | $k_{a2}$ (1/RUs)* | | | | |
| | $k_{d2}$ (1/s) | | | | |
| | $K_{A1}$ (1/M) | | | | |
| | $K_A$ (1/M)** | $6.56 \times 10^{5}$ | $1.74 \times 10^{6}$ | | |
| | $K_{D1}$ (M) | | | | |
| | $K_D$ (M) | $3.7 \times 10^{-6}$ | $1.3 \times 10^{-6}$ | | |
| | $\chi^2$[%] | 0.31 | 0.54 | | |
| $AbrB_{BA}$:$AbbA_{BA}$ | $k_{a1}$ (1/Ms) | $1.30 \times 10^{4}$ | $1.26 \times 10^{4}$ | $8.22 \times 10^{4}$ | $1.32 \times 10^{5}$ |
| | $k_{d1}$ (1/s) | $1.15 \times 10^{-2}$ | $1.08 \times 10^{-2}$ | $2.54 \times 10^{-3}$ | $9.55 \times 10^{-2}$ |
| | $k_{a2}$ (1/RUs)* | | | $2.78 \times 10^{-2}$ | $1.78 \times 10^{-2}$ |
| | $k_{d2}$ (1/s) | | | $2.25 \times 10^{-2}$ | $1.58 \times 10^{-3}$ |
| | $K_{A1}$ (1/M) | | | $3.34 \times 10^{7}$ | $1.33 \times 10^{6}$ |
| | $K_A$ (1/M)** | $1.32 \times 10^{6}$ | $1.23 \times 10^{6}$ | $3.18 \times 10^{7}$ | $1.5 \times 10^{7}$ |
| | $K_{D1}$ (M) | | | $3.45 \times 10^{-8}$ | $7.91 \times 10^{-7}$ |
| | $K_D$ (M) | $1.14 \times 10^{-6}$ | $1.24 \times 10^{-6}$ | $3.14 \times 10^{-8}$ | $6.89 \times 10^{-8}$ |
| | $\chi^2$[%] | 0.38 | 1.57 | 0.65 | 0.19 |

**SM4.** Dissociation of bound AbbA analytes from AbrB ligands expressed as percentage. Dissociation period followed after the end of injection. The dissociation started as of 140 seconds, this time point was set 100 %: (AbbA bound (RU)/ AbbA bound (RU) at t=140 sec) x 100 %. Dissociation phase observed with different AbbA concentrations (0.25 µM, 0.5 µM, 0.75 µM and 1.0 µM). BS = *B. subtilis* 168, BA = *B. amyloliquefaciens* FZB45.

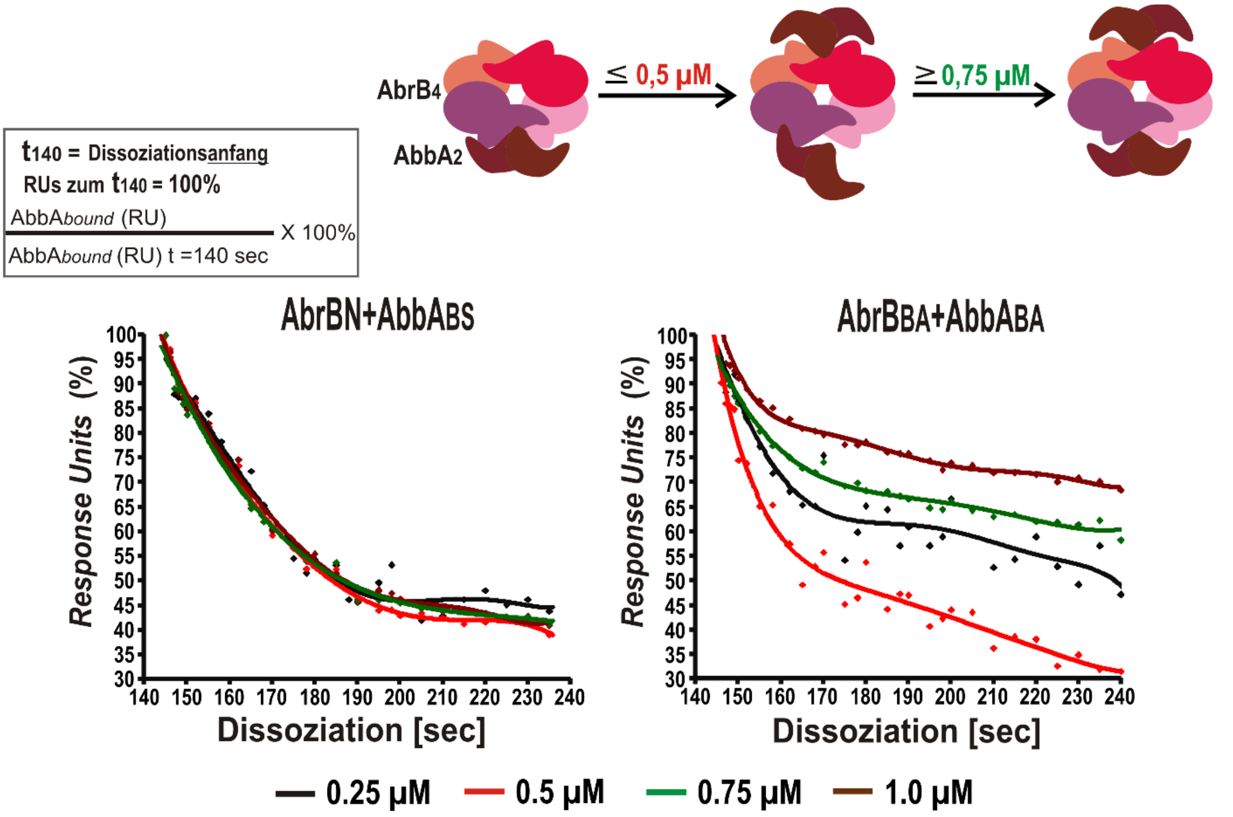

**SM5**.Important parameters derived from scattering data. GASBOR reads in output files of GNOM including exctracted parameters (such as $R_g$, $D_{max}$) and particle symmetry. $R_g$ is radius of gyration, $D_{max}$ is maximum particle diameter. Output file from GASBOR contains information for number of C-α atoms and H atoms meaning positions of dummy residues and dummy bound waters. The number of dummy residues should be equal to number of amino acid residues in the protein. Theoretical molecular mass correspond to polymer form of proteins (AbrB tetramer and AbbA dimer).

| **parameters** | **AbrB** | **AbbA** |
|---|---|---|
| momentum transfer range s [$nm^{-1}$] | 0.14 to 2.59 | 1.21 to 3.42 |
| $D_{max}$ [nm] | 13.5 | 13.5 |
| $R_g$ [nm] | 3.86 | 2.83 |
| $s_{max}$ [1/Å] | 0.259 | 0.342 |
| particle symmetry | P22 | P2 |
| dummy $H_2O$ molecules | 378 | 90 |
| dummy residues | 460 | 130 |
| Molecular mass, theory (kDa) | 50.56 | 15.78 |
| Molecular mass, PRIMUS (kDa) | 40 | 16 |

**SM6**. Scattering curves for AbrB (A) and AbbA (B). Dilute aqueous solutions of proteins give rise to an isotropic scattering intensity (I), which depends on the modulus of the momentum transfer (s, $nm^{-1}$). Following subtraction of the solvent (dialysis buffer) scattering, the background corrected intensity I(s) is proportional to the scattering of a single particle averaged over al orientations. Data from both protein concentrations (2.5 mg/ml and 5 mg/ml) were merged to give the final data file. The final curve (averaged, subtracted, merged) was fitted and evaluated to provide the several overall important parameters.

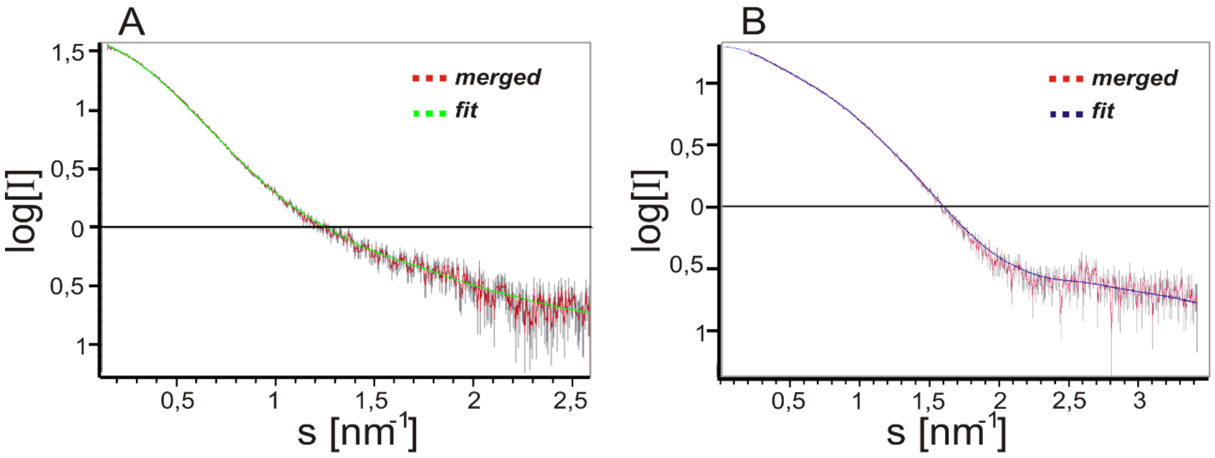

**SM7.** Low resolution models of the AbbA dimer calculated from the scattering data assuming P2 symmetry. Three views rotated about longitudinal axis clarifying the differently orientated protein sidewise arms.

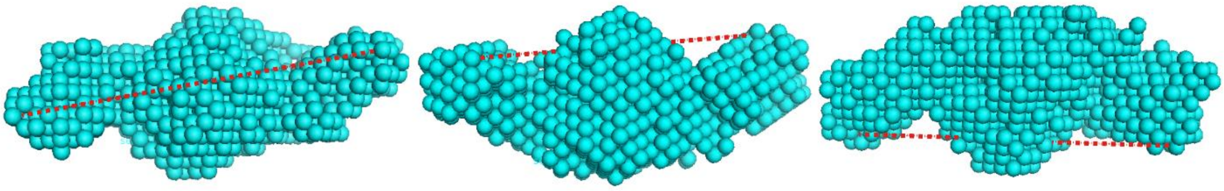
